# High-throughput screening and structure-guided design of small molecules enable modulation of SAL-PAP stress signaling

**DOI:** 10.1101/2025.10.20.683270

**Authors:** Nay Chi Khin, Melanie Carmody, Brett D. Schwartz, Michael Dlugosch, Suyan Yee, Marten Moore, Michael Gardiner, Li Lynn Tan, Colin Jackson, Ben Corry, Lara R. Malins, Barry Pogson, Kai Xun Chan

## Abstract

Chloroplasts sense environmental stress and activate chloroplast-to-nucleus retrograde signalling, reprogramming nuclear gene expression to drive plant acclimation. One such pathway is regulated by the chloroplastic phosphatase, SAL, which hydrolyses the nucleotide signal 3’-phosphoadenosine 5’-phosphate (PAP). In *Arabidopsis thaliana*, genetic loss of AtSAL1 elevates PAP and enhances stress tolerance but causes pleiotropic growth defects, complicating the interpretation of the role of PAP in cellular signalling. To uncouple stress signalling from genetic pleiotropy, we conducted a high-throughput *in vitro* screen of 13,000 small molecules and identified **V20,** a competitive inhibitor of AtSAL1 with three-fold greater potency than the known Li⁺ inhibitor. Structural analoguing of **V20** and biochemical assays defined key pharmacophore features required for inhibition, including an ortho-halogenated aromatic ring, hydrogen-bond donor capacity at the triazole carboxamide, and electron-withdrawing substitutions that enhance π–π stacking with aromatic residues. Accelerated molecular dynamics simulations using a new high-resolution crystal structure revealed two previously uncharacterised **V20** binding pockets adjacent to the catalytic site. **V20** binding induces conformational changes which restrict substrate access to the catalytic site. Exogenous application of **V20** to *Arabidopsis* led to increased PAP accumulation, activated PAP-responsive gene expression and enhanced oxidative tolerance, demonstrating cellular uptake and *in vivo* efficacy in the energy organelles, chloroplasts and mitochondria, where AtSAL1 is localised. Collectively, our findings reveal new insights into the regulatory domains of SAL enzymatic activity for control of PAP-mediated signalling and establish a proof-of-concept for targeted chemical modulation of SAL activity, which offers novel strategies to selectively manipulate chloroplast-to-nucleus retrograde signalling in plants.

**Significance Statement:** We applied drug discovery strategies to test 13,000 small molecules for inhibition of AtSAL1, a key regulator of plant stress signalling from chloroplasts. Using biochemical assays, chemical analogues and molecular dynamic simulations, we defined the structural-activity relationship of a lead compound, **V20**, and elucidated its inhibitory mechanism. Application of **V20** to plant leaves elevated the levels of AtSAL1’s natural substrate, PAP (a chloroplast stress signalling molecule), activated stress-responsive genes and improved oxidative stress tolerance in Arabidopsis. This on-demand activation of SAL-PAP chloroplast signalling avoids pleotropic effects observed in loss-of-function mutants. Our approach provides a new strategy to study chloroplast-mediated cellular signaling pathways that are crucial for plant acclimation to stressful environmental conditions such as excessive sunlight.

## Introduction

When plants experience oxidative stress such as during exposure to excessive sunlight, chloroplasts initiate retrograde signalling to activate nuclear-encoded stress responses, enabling physiological and acclimatory adjustments (1). One key retrograde signal is 3’-phosphoadenosine-5’-phosphate (PAP), a byproduct of sulfur metabolism that accumulates under oxidative stress (2, 3). The chloroplast nucleotide phosphatase, SAL, plays a crucial role in PAP turnover under non-stress conditions, thereby maintaining *in vivo* PAP concentration at a basal level. SAL is widely conserved across species, from bacteria to humans (4–7), and is well-characterised in the model plant *Arabidopsis thaliana* as a regulator of PAP-mediated retrograde signalling (8–13). During abiotic stresses such as drought and excessive light, redox-dependent inactivation of SAL leads to PAP accumulation (14). PAP can move between chloroplasts and the cytosol via PAP/PAPS transporters (PAPSTs) (15) regulate gene expression via inhibition of exoribonucleases (XRNs) (2, 16) leading to the activation of drought-, and high light-inducible genes (14, 17, 18).

In *Arabidopsis*, loss of AtSAL1 function leads to PAP over-accumulation, enhancing drought and high light tolerance compared to wild-type (WT) plants (2, 3). However, constitutive PAP elevation in *atsal1* mutants results in pleiotropic effects, impacting hormonal homeostasis, growth, and development (9, 19–21). Consequently, the ‘true’ impacts of PAP accumulation on cellular signalling are likely to be partially masked by confounding factors from these pleiotropic effects.

A key challenge in modulating PAP levels for stress tolerance without adverse effects on plant growth is that SAL is a stable protein with a low turnover rate (22) and that SAL is an efficient enzyme, as indicated by its kinetic parameters against PAP (14). Effective PAP accumulation *in vivo* requires near-complete loss of SAL activity to trigger transcriptomic and physiological changes. Previous efforts to modulate PAP levels for stress tolerance without adverse effects on plant growth through genetic manipulation (RNAi silencing) have been unsuccessful (22). A recent paper attempted to prime PAP accumulation by selective knockout of SAL homeologues in wheat, which resulted in improved stress resilience and water productivity and better yield stability (23). However, single knockouts of two different wheat SAL homeologues had opposite impacts on these phenotypes (23), demonstrating the unpredictability in genetic manipulation of SAL.

Exogenous PAP application, while effective on isolated tissues like epidermal peels (24), has no effect on whole plants likely due to the inability of PAP to cross the leaf cuticle (2). However, given that SAL biochemical activity can be post-translationally regulated by hydrogen peroxide and glutathione (14) as well as by monovalent ions such as Li^+^ (6), modulation of SAL activity using small molecules could be a viable alternative.

High-throughput screening (HTS) of chemical libraries is used in modern drug discovery, enabling the identification of “hit” compounds that can modulate biological targets. These small molecules can be applied in a reversible, dose-dependent, and conditional manner, offering precise temporal control over biological systems (25). In mammalian systems, HTS has been widely employed for drug discovery across a range of human diseases (26). In plants, chemical screening strategies have proven to be efficient tools for uncovering compounds involved in diverse biological processes (27, 28).

There are two primary HTS approaches: *in vitro* biochemical assays for target-based drug discovery and cell-based or *in vivo* assays for phenotype-based drug discovery (29). Phenotypic, or forward, screening aims to dissect biological processes by identifying bioactive molecules that induce observable phenotypes. This approach allows for the simultaneous monitoring of multiple biochemical and morphological parameters, enabling the discovery of broad phenotypic responses and previously uncharacterised biological mechanisms (30). For example, phenotype-based screens in plants have identified small molecules that enhance selenium accumulation or confer tolerance to selenate toxicity by targeting β-glucosidase in *Arabidopsis* (31). In herbicide discovery, phenotypic screening has proven effective in identifying inhibitors of essential biosynthetic enzymes, such as those involved in branched-chain amino acid and fatty acid synthesis (32, 33).

In contrast, target-based reverse screening (employed in this study) focuses on discovering compounds that modulate the activity of known protein targets, using assays that measure enzymatic activity, protein–protein interactions, or transcription factor binding (34–36). In human drug discovery programs, the majority of approved small molecule drugs have emerged from phenotypic rather than target-based approaches (37). Similarly, in plants, there are relatively few examples of target-based screening, likely due to challenges such as compound delivery, detoxification, and increasingly stringent regulations (38). However, when the target is physiologically well-characterised, as is the case for SAL, target-based screening can enable the development of highly specific chemical tools to precisely modulate targeted biochemical pathways and/or signalling networks.

In this study, we utilised a reverse chemical screening approach to deactivate the SAL1 enzyme, with the goal of enabling controlled elevation of PAP levels and activation of PAP-mediated signalling in plants. Our approach involved HTS of a chemical library to identify small molecules that inhibit recombinant AtSAL1 protein with higher specificity and potency than the current known monovalent ion inhibitor, Li^+^. Through enzymatic assays, chemical synthesis of small molecule analogues, and *in silico* molecular dynamics (MD) simulations, we identified novel compounds and mapped their interactions with potential regulatory regions of AtSAL1. Lastly, we demonstrated the *in vivo* activity of a novel AtSAL1 inhibitor, **V20**, which elevates PAP levels, activates PAP-induced stress response genes and enhances tolerance to oxidative stress when applied to *Arabidopsis* plants.

## Results

### Screening of chemical inhibitors of AtSAL1 enzymatic activity

To identify small molecules with inhibitory potency against recombinant *Arabidopsis* SAL1 (AtSAL1) enzyme *in vitro*, we developed a high-throughput screen (HTS) based on a colorimetric enzyme assay with malachite green, which detects the production of inorganic phosphate (P_i_) from AtSAL1-mediated PAP degradation (Figure S1A) (39). In the reaction for positive control, adding malachite green to AtSAL1 preincubated with PAP produced a dark green phospho-molybdate complex, reflected by a high absorbance at 623 nm (A_623_). In contrast, the inhibition control (AtSAL1 with 0.5 mM LiCl and PAP) and the negative control (AtSAL1 only without PAP) exhibited significantly reduced phospho-molybdate complex formation, resulting in A_623_ values that were 4.6-fold and 7.2-fold lower than the positive control, respectively (Figure S1B, C). We calculated the Z’ factor, a measure of assay robustness for high-throughput screening (40), as 0.64 (Figure S1D), indicating that this assay is sensitive and reliable for detecting AtSAL1 enzymatic activity in the presence of potential small molecule inhibitors.

Using this colorimetric assay, we conducted a high-throughput screening of 13,000 compounds from the Chembridge DIVERSET® library (Figure 1A). The primary screen identified 388 compounds as putative AtSAL1 inhibitors (Figure 1B). To exclude false positives, these compounds underwent secondary screening with high-performance liquid chromatography (HPLC), which specifically quantified AMP instead of P_i_ from AtSAL1-mediated PAP degradation (Figure S1E, F). This refined the list to 97 compounds, collectively designated as **V1**–**V97** (Figure 1C). 81% of the inhibitors (79 compounds) shared one of eight common core scaffolds, designated as Groups 1-8, while the remaining compounds were classified as Group 9 (Figure S2).

**Figure 1.**
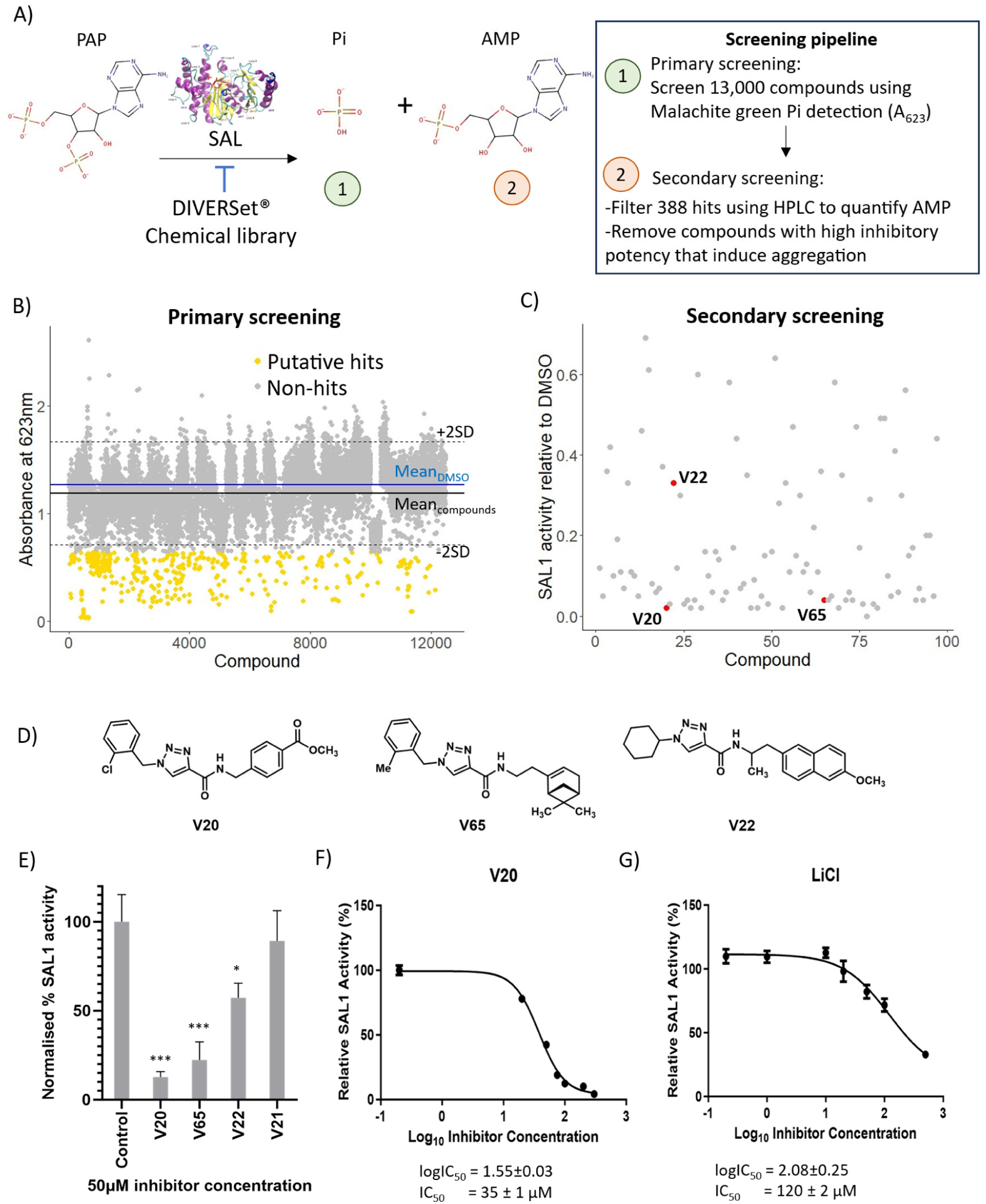
High-throughput screening identifies small molecule inhibitors of Arabidopsis SAL (AtSAL1). **(A)** Schematic overview of the two-stage screening pipeline. In the primary screen, ∼13,000 small molecules were tested for inhibition of AtSAL1 phosphatase activity using a malachite green-based colorimetric assay to detect released inorganic phosphate (Pi) at 623 nm (A₆₂₃). Compounds that significantly reduced Pi release were considered putative hits. In the secondary screen, 388 of these hits were re-evaluated using an HPLC-based AMP quantification assay to confirm inhibitory activity and reduce false positives caused by compound aggregation. **(B)** Distribution of compounds in the primary screen based on absorbance at 623 nm. The mean ± 2 standard deviations (SD) of DMSO controls is shown in blue. Compounds with absorbance values below −2 SD from the mean (yellow) were classified as putative hits (388 total). **(C)** Secondary screening of 388 primary hits based on HPLC-quantified AMP production relative to DMSO control. A subset of compounds identified as non-aggregators from detergent sensitivity assay is depicted in red and selected for further study (Figure S3b). **(D)** Chemical structures of three non-aggregating lead compounds, which share a common Group 2 scaffold (triazole carboxamide) and were selected for further analysis. **(E)** AtSAL1 inhibitory potency of Group 2 core structure compounds at 50 µM, determined by HPLC-based AMP quantification. Data shown relative to DMSO control (n=3). Asterisks indicate statistical significance compared to control: P < 0.05 (*); P < 0.01 (**); P < 0.001 (***). **(F, G)** Dose–response curves showing IC₅₀ values of (F) **V20** and (G) LiCl, a known SAL inhibitor (n=3). The error bars indicate standard errors.

Colloidal compound aggregation, a common phenomenon among HTS hits, can cause local protein denaturation and apparent inhibition of enzyme activity (41, 42). To assess whether aggregation contributed to the observed inhibition, we tested the most potent compounds from Groups 1–9 in the absence and presence of the non-ionic surfactant Triton X-100 (TTX-100), which disrupts aggregate formation. All compounds, except those from Group 2, exhibited either complete or partial loss of inhibitory potency in the presence of TTX-100, indicating their activity likely relies on aggregation (Supplemental Figure 3B). In contrast, three Group 2 compounds (**V20**, **V65** and **V22**), along with the known SAL inhibitor, LiCl, showed minimal differences in inhibitory potency with or without TTX-100, suggesting their inhibition does not rely on aggregate formation. Consequently, we identified these three compounds from Group 2 as genuine inhibitors of AtSAL1 *in vitro* (Figure 1C, D), with **V20** being the most potent (Figure 1E). The IC₅₀ of **V20** was 35.3 µM, representing a 3.4-fold increase in potency compared with the known inhibitor LiCl, which has an IC₅₀ of 120.3 µM (Figure 1F, G).

### Chemical analoguing reveals key structural determinants of V20 required for its inhibitory activity

All three AtSAL1 inhibitors identified shared a 1*H*-1,2,3-triazole-4-carboxamido motif, with **V22** having substantially lower inhibitory potency compared to **V20** and **V65** (Figure 1D, E). We therefore sought to identify the crucial pharmacophoric elements required for **V20** to inhibit AtSAL1 activity. We rationally modified the functional groups (i.e., substitution, addition, and deletion) of **V20** through a series of chemical analogues. Accordingly, **V20** was synthesised along with a library of an additional 26 analogues, which were tested for their inhibitory potencies. Structural modifications were introduced in three distinct regions of **V20**, designated as Rings A, B, and C (Figure 2A), with particular emphasis placed on modifications to Ring C. We performed *in vitro* assays of AtSAL1 activity in the presence of each analogue individually, both in the presence or absence of detergent, and confirmed that all these compounds function by non-aggregation mechanisms (Figure 2B, C).

**Figure 2.**
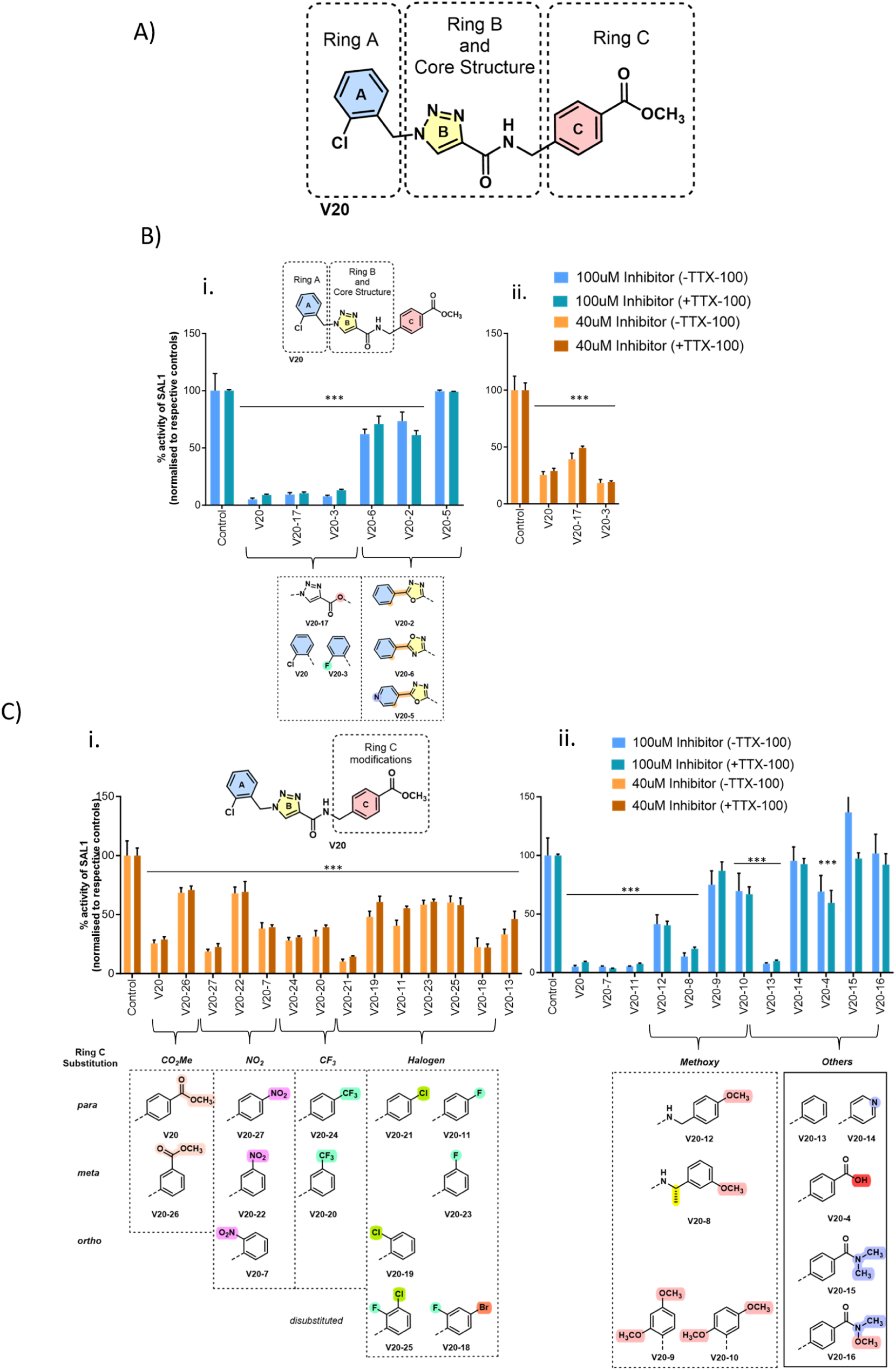
Analogue profiling of the V20 lead compound reveals key pharmacophore features required for AtSAL1 inhibition. **(A)** Chemical structure of **V20** showing its modular components: Ring A (2-chlorophenyl), Ring B (1*H*-1,2,3-triazole carboxamide), and Ring C *(p-*carbomethoxyphenyl) moieties. **(B, C)** Inhibitory activity of **V20** analogues at 100 µM (blue bars) and 40 µM (orange bars) concentrations, assessed using an HPLC-based AMP assay at 85 µM PAP, in the presence and absence of Triton X-100 (TTX-100). AtSAL1 activity is normalized to the DMSO control (n=3-6). Analogues based on lead compound **V20**. Only the varied regions of the molecules are shown and modifications are highlighted in colour; conserved core structures are omitted for clarity. Substituent changes are highlighted to illustrate structure–activity relationship (SAR) trends. **(B)** Analogues with modifications to Rings A and B **(C)** Analogues with modifications to Ring C. Substituents are colour-coded by functional group. The Tukey’s multiple comparison test was performed between the control and each inhibitor treatment with significance represented by asterisks (*P ≤ 0.05, **P ≤ 0.01, ***P ≤ 0.001). Error bars indicate standard deviations.

Modifications made to the Ring A chlorobenzene moiety revealed that substituting the electron-withdrawing chlorine (Cl) atom in the chlorobenzene ring with a smaller, more electronegative halide such as fluorine (F) (**V20-3**) enhanced inhibition by approximately 10% (Figure 2Bii). Conversely, simultaneous removal of the chlorine from Ring A and replacement of the triazole core structure with oxadiazoles (**V20-2**, **V20-5**, **V20-6**) drastically diminished inhibitory activity (Figure 2Bi). Potency was also reduced by approximately 20% when the amide (N–H) attached to Ring B was substituted with an ester (**V20-17**). This is likely due to the loss of hydrogen bond donor capacity of the amide.

Substitution of the methyl ester on Ring C of **V20** with various electron-withdrawing groups at the *para* position resulted in variable effects on potency, which generally followed the trend: carboxylic acid (COOH, **V20-4**) < fluoro (**V20-11**) < CF₃ (**V20-24**) < NO₂ (**V20-27**) < chloro (**V20-21**). Interestingly, the absence of a substituent at this position (**V20-13**) did not significantly reduce potency (Figure 2C). The 4-bromo-2-fluoro analogue (**V20-18**) was 25% more potent than **V20** at 40 μM inhibitor concentration. However, substituting a pyridine (**V20-14**) in place of carbon at C-4 of Ring C, or replacing the methyl ester with methoxy groups (**V20-12**, **V20-8**, **V20-9**, **V20-10**) at various positions around the ring reduced potency. Electron-withdrawing groups at the *para* and *ortho* positions were tolerated more than those at the *meta* position, particularly for ester and nitro substituents (e.g. **V20-26**, **V20-22**) on the benzene ring. The presence of a *para*-substituted carboxylic acid (**V20-4**) on Ring C or tertiary amides (**V20-15**, **V20-16**) resulted in near-complete loss of inhibitory activity.

In summary, analysis of the 27 analogues in our **V20** library revealed that inhibitory activity is driven by a combination of key features: electron-withdrawing substituents such as an *ortho*-substituted halogen on Ring A, the hydrogen bond donor capability associated with the amide appended to the Ring B and π–π stacking interactions of the triazole core (Ring B), and electron-withdrawing groups in the *para* position of Ring C. Among Ring C variants, the *para*-chloro analogue (**V20-21**) was the most effective, with *para*-nitro (**V20-27**) also showing improved inhibition of AtSAL1, relative to other substitutions. Future work will focus on further exploration of Ring A substitutions to optimise potency.

### Accelerated molecular dynamics for determination of AtSAL1 inhibitor binding sites

To overcome the limitations of our previous low-resolution and twinned crystal structure of AtSAL1 (14), we pursued new crystallisation conditions to obtain an improved atomic model. This effort yielded a new, untwinned crystal form in the *P*6₁22 space group that diffracted to a higher resolution of 2.60 Å. The resulting electron density map was significantly clearer, which enabled the unambiguous modelling of previously disordered surface loops and side chains, providing a more complete and accurate view of the enzyme’s structure. We then sought to identify the putative binding site(s) on the new AtSAL1 structure and mechanism(s) of action for **V20** through accelerated molecular dynamics (aMD) simulations.

To establish the robustness of this approach we first examined the binding of PAP, the natural substrate, to AtSAL1 using 6 µs aMD (Figure S4A). In replicate 1, PAP binds to the catalytic site with its 3’ phosphate group initially interacting with the Mg²⁺ before rotating to a more stable orientation, where the 5′ phosphate group forms a persistent interaction with Mg²⁺ for the remainder of the simulation (Figure S4B). Binding is initiated by interactions with positive charged K36 and K39 on Loop 1 and subsequently guided into the active site by residues such as K140 and R144 on Loop 5, as well as D258, which are positioned above the catalytic pocket, acting as a gate into the active site (Figure S4B). These observations corroborate the previously proposed regulatory role of Loop 1 in SAL enzymatic activity (14) and are further supported by mutagenesis data showing that replacement of PAP-interacting residues on Loop 1 (D32, K36) and Loop 5 (D146, R144) with alanine leads to partial or complete loss of AtSAL1 enzyme activity (Figure S5); as well as the crystal structure of the homolog ScHAL2 bound to PAP (43, 44). Together with mutagenesis and biochemical assays, these results validate the reliability of aMD simulations for identifying potential binding sites and elucidating the mechanisms of action for novel small molecules that modulate AtSAL1 activity.

Applying the aMD approach to the **V20** allowed us to identify stable binding sites for the inhibitor at two distinct sites on the protein (Replicate 1 cluster 2 and Replicate 5 cluster 1), which we named as candidate binding site 1 and 2 (CBS1 and CBS2) respectively (Figure 3A, B; Supplementary Figure 6). Both CBS1 and CBS2 were located close to the active site, with CBS1 situated behind the catalytic site and CBS2 at the base of Loop 1. At CBS1, the binding of **V20** to the protein is stabilized by hydrophobic interactions with Ring A (P171, L172, A173, V216), parallel π-π stacking and CH_3_¼p interactions (F192 and A265 to Ring C, respectively) and hydrogen bonding between the polar core structure of V20 and Y148 and the backbone of the protein (L266, S267, R268) (Figure 3C, Supplementary Figure 7A). Similarly, at CBS2, the most favourable protein-ligand interactions come from the hydrogen bonds formed by the protein backbone (G48, S49, Q50, A51, V52, V53) and the side chains of Q25 and S49 with the core scaffold and Ring B of **V20**, followed by π-π stacking interactions between the aromatic rings of Y148 and Ring C and hydrophobic interactions (V52, A51 and methylene group of D84) with Ring A (Figure 3D, Supplementary Figure 7B).

**Figure 3.**
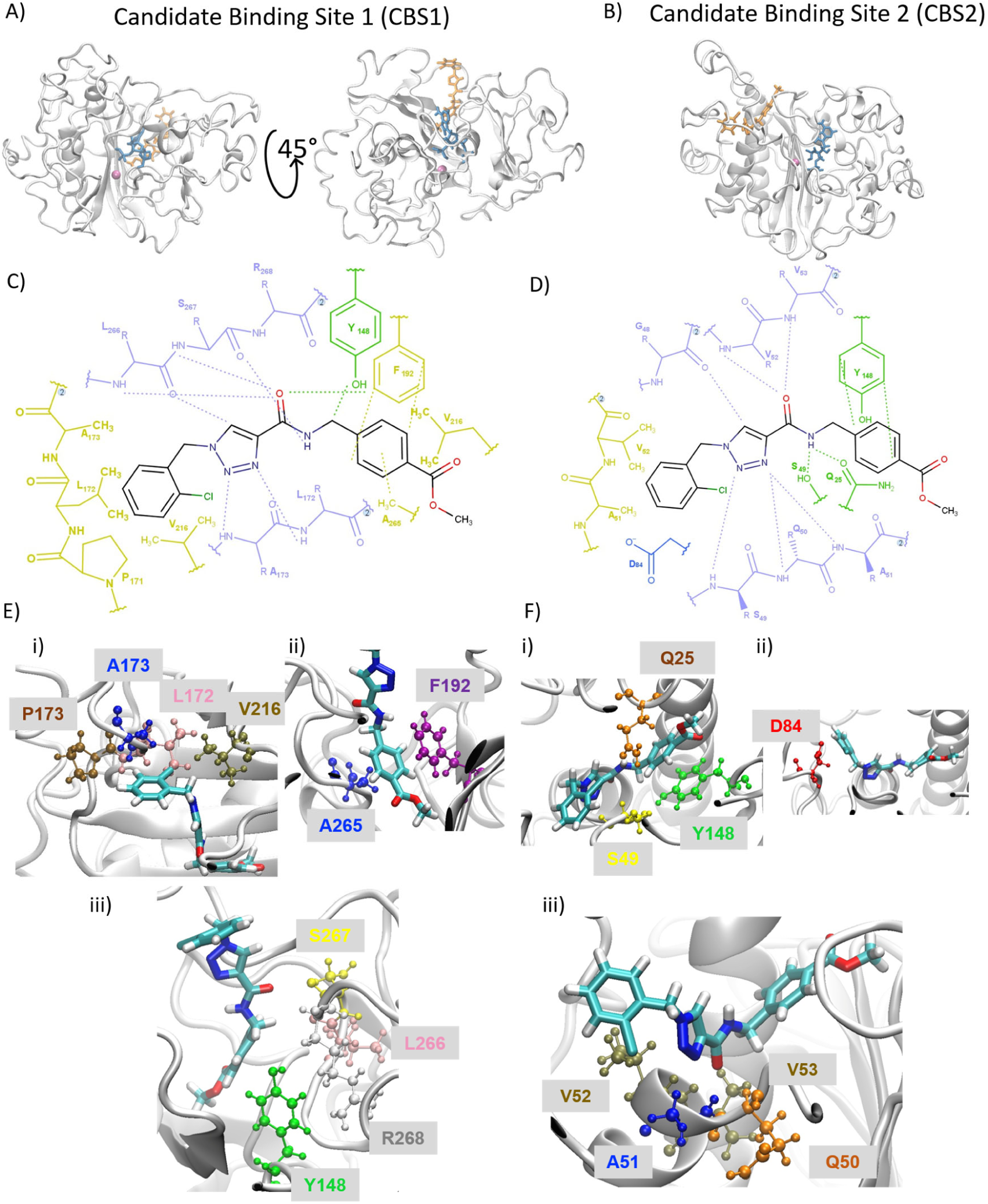
Accelerated molecular dynamics (aMD) simulations reveal two stable V20 binding sites on AtSAL1. **(A, B)** Representative **V20** conformation from aMD simulations showing two stable binding clusters (Figure S6) on the AtSAL1 protein, designated as candidate binding sites (CBS): **(A)** CBS1 and **(B)** CBS2. AtSAL1 is shown in cartoon representation; the **V20** ligand is displayed in an orange licorice model, and the PAP molecule (to indicate the active site) is shown in blue. **(C, D)** Predicted interactions between **V20** and AtSAL1 residues with the most favourable interaction energies (see Figure S7) at (C) CBS1 and (D) CBS2. Residues are shown as either side chains or backbones, color-coded by chemical property: positively charged (red), negatively charged (blue), polar (green), non-polar (yellow), and backbone (purple). Certain residues (e.g., S49, V216) appear twice in the 2D depiction because the ligand adopts multiple contact points in 3D space during simulation. **(E, F)** Close-up views of key atomic interactions between **V20** and surrounding AtSAL1 residues at **(E)** CBS1 and **(F)** CBS2. Interacting residues are boxed and labelled by residue colours.

**V20** maintained 93-99.7% ligand residency at CBS1 and CBS2 with minimal ligand-ligand or transient interactions, similar to PAP binding at the active site (99.3% ligand residency) (Figure S7C). As an additional control, we performed aMD with two aggregator compounds, **V30** and **V01**, and observed no stable binding to AtSAL1 as expected (Figure S8).

We then performed biochemical studies to validate the candidate binding sites found using aMD. Michaelis-Menten kinetics of AtSAL1 in presence of **V20** suggest competitive inhibition of AtSAL1 activity (Figure S9), as K_m_ showed a substantial shift by approx. 60-fold whereas *k_c_*_at_ was only marginally altered with increasing inhibitor concentration. This indicates that **V20** inhibits AtSAL1 by competing with substrate binding to the active site, lending support to CBS1 that resides adjacent to the active site. We then tested the ability of **V20** to inhibit activity of AtSAL1 proteins carrying individual point mutations in residues predicted by aMD to be involved in PAP binding and catalysis to uncover how changes in PAP binding might modulate **V20**’s inhibitory potency (Figure S10). As expected, some of these mutations in loop 1 (Y47A and S35A) and elsewhere on the protein (F103A, R144A, V52A) caused drastically diminished enzymatic activity; while other mutations (S185A and the gating residues K140A and D258A) strongly enhanced AtSAL1 catalytic activity of PAP, suggesting that these residues play a regulatory role and may be rate-limiting the enzyme’s activity (Figure S5B). Of the 11 mutants tested (excluding those with complete or near-complete loss of activity), four mutations (K140A, D258A, S35A, and D146A) substantially affected **V20**’s inhibitory potency (Figure S10A). These residues do not directly interact with **V20** at CBS1 and CBS2 binding sites and may indirectly influence the potency of **V20** by altering the conformational dynamics of AtSAL1 upon ligand binding, thereby modulating the extent of inhibition (Figure S10B). K140 and D258 are predicted to undergo conformational changes required for substrate binding, potentially influencing the accessibility or stability of the active site (Figure S4). At CBS1, K140A, D258A, and D146A likely disrupt charged interactions that help anchor Loop 1, affecting its flexibility and conformation. At CBS2, K140A and D258A appear to participate in a charged residue network that modulates access to the catalytic core and influences Loop 1 mobility. Collectively, these biochemical results support the aMD predictions and establish **V20** as a bona fide AtSAL1 inhibitor, that may engage at least two functionally relevant and conformationally sensitive sites on the protein.

### V20 functions as a AtSAL1 inhibitor in planta to confer oxidative stress tolerance and mediate PAP-induced transcriptomic changes

To determine whether **V20** can function as a AtSAL1 inhibitor *in vivo*, we first fed either mock solution, PAP alone, **V20** alone, or **V20** and PAP to isolated *Arabidopsis* protoplasts, which are devoid of cell walls and consequently more permeable to exogenous metabolites (45). We then quantified PAP levels in these protoplasts 1 h post-treatment. Protoplasts treated with either mock or PAP alone had similar levels of PAP, consistent with the high PAP degradation activity of endogenous SAL (14, 22). Similarly, protoplasts treated with **V20** alone did not show significant PAP accumulation, likely due to the low rates of endogenous PAP biosynthesis under unstressed conditions (22). In contrast, protoplasts co-treated with both **V20** and PAP exhibited significantly higher PAP accumulation compared to those treated with PAP alone (Figure 4A), suggesting that **V20** successfully inhibited AtSAL1 catalytic activity *in vivo* thus allowing the exogenous PAP to accumulate.

**Figure 4.**
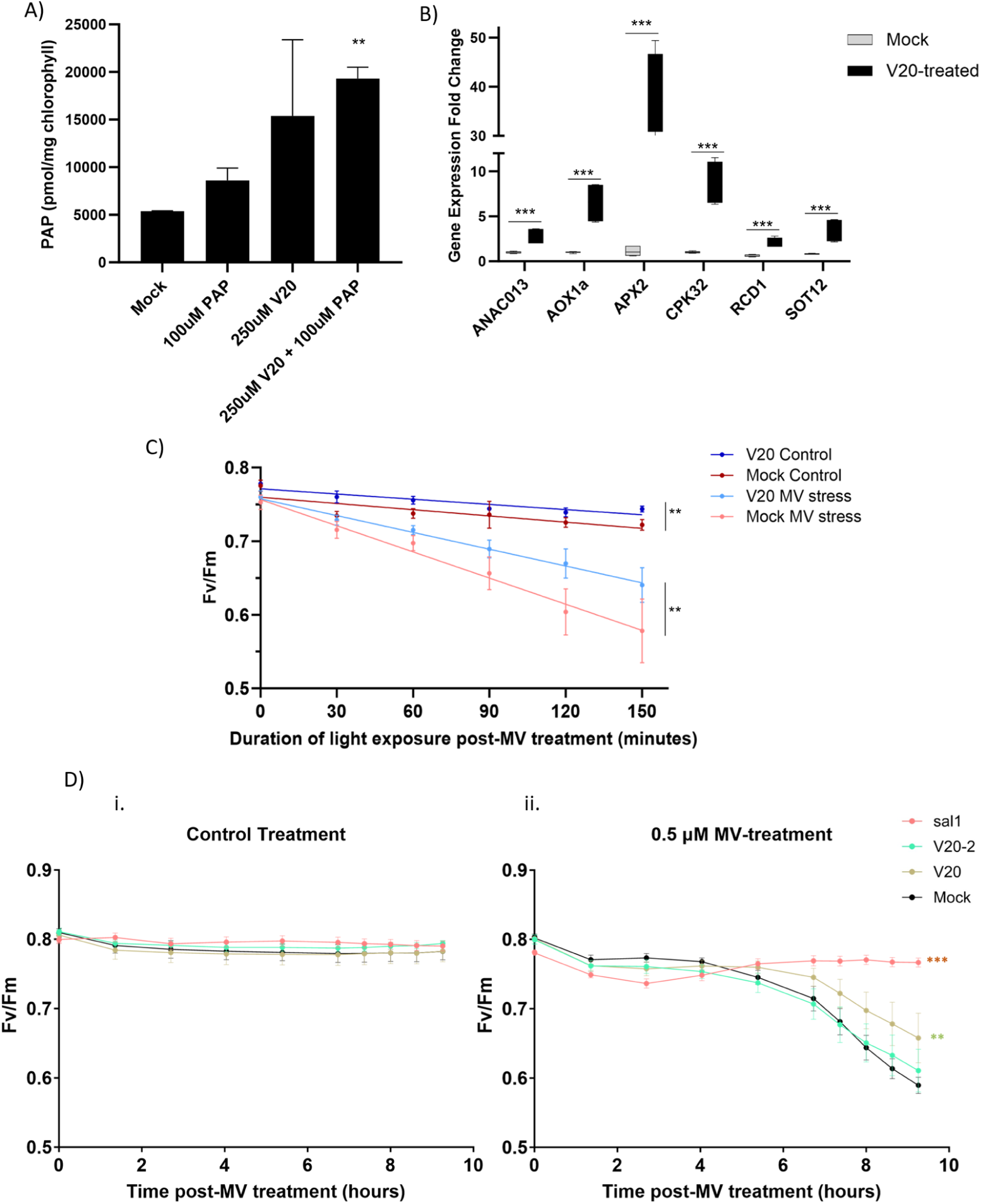
V20 functions as a AtSAL1 inhibitor in planta to enhance oxidative stress tolerance and activate PAP-responsive gene expression. **(A)** Quantification of PAP levels in Arabidopsis protoplasts 1 hour after treatment with PAP (50 µM) alone or in combination with **V20** (500 µM) (n = 2-3). **(B)** Transcript levels of known PAP-responsive genes (*Ascorbate Peroxidase 2 (APX2), Alternative Oxidase 1a (AOX1a), Calcium Dependent Protein Kinase 32 (CPK32), Arabidopsis NAC domain containing protein 13 (ANAC013) and Sulfotransferase 12 (SOT12)*) measured in whole plants treated with mock solution or **V20** (500 µM) (n = 3). **(C)** Time course of Fv/Fm in whole Arabidopsis plants treated with mock or **V20** (500 µM), with and without methyl viologen (MV) application. Measurements began following exposure to high light (n = 5–6). **(D)** Time course of Fv/Fm in Arabidopsis Col-0 and *atsal1* leaf discs infiltrated with mock, **V20** (500 µM), or the weak analogue **V20-2** (500 µM), under conditions (i) without and (ii) with MV (0.5 µM) treatment and light exposure (n = 4–6). The error bars indicate SEM. Asterisks denote significant differences between mock and inhibitor-treated samples (*P < 0.05, **P < 0.01, ***P < 0.001; Student’s t-test or ANOVA as appropriate).

Next, we tested if **V20** application on whole plants could activate PAP-mediated chloroplast signalling by measuring the expression of stress-induced and ROS-responsive genes known to be activated by PAP signalling [*Alternative Oxidase 1a (AOX1a), Ascorbate Peroxidase 2 (APX2), Calcium Dependent Protein Kinase 32 (CPK32), Arabidopsis NAC domain containing protein 13 (ANAC013), Radical Cell Death Induced 1 (RCD1), and Sulfotransferase 12 (SOT12)*] (Figure 4B).

**V20** treatment significantly increased the expression of these PAP signalling genes compared to mock treatment (Figure 4B), suggesting the activation of PAP signalling through inhibition of AtSAL1 by **V20** application.

We then tested whether the up-regulation of stress responsive genes by **V20** could alter plant oxidative stress tolerance by challenging mock- and **V20**-treated plants with either Control (water) or methyl viologen (MV) in *Arabidopsis* whole plants and leaf discs. MV is an electron mediator at photosystem I that promotes reactive oxygen species (ROS) accumulation upon exposure to light, causing photoinhibition as shown by a decrease in the PSII quantum yield, Fv/Fm. We found that while both mock- and **V20**-treated plants had similar Fv/Fm values, **V20**-treated plants had enhanced oxidative stress tolerance as evidenced by a slower decrease in Fv/Fm compared to mock-treated whole plants after MV challenge (Figure 4C). Finally, we tested whether **V20** was genuinely responsible for the increased oxidative stress tolerance observed in Figure 4C by comparing its effect on Fv/Fm in wild type Col-0 *Arabidopsis* under MV treatment to that of **V20-2** (weak **V20** analogue, Figure 2C) and constitutive PAP accumulation in the *atsal1* mutant. We observed that leaf discs without MV, regardless of mock or inhibitor treatment, maintained Fv/Fm values around 0.8 over a 10-hour period, indicating the lack of toxicity in leaf disc preparation and **V20**/**V20-2** treatment over this period (Figure 4D). Under MV treatment, Fv/Fm progressively and significantly declined in mock-treated wild-type (WT) leaf discs, while *atsal1* leaf discs maintained Fv/Fm as expected. Notably, WT leaf discs treated with **V20** exhibited significantly higher Fv/Fm compared to leaf discs treated with mock solution (Figure 4D). Consistent with its weaker inhibitory potency against AtSAL1 *in vitro* (Figure 2Bi), under MV challenge the leaf discs treated with **V20-2** had significantly lower Fv/Fm compared to **V20**-treated leaf discs, and similar Fv/Fm to mock-treated leaf discs (Figure 4D).

Taken together, these findings indicate **V20** effectively inhibits AtSAL1 *in vivo* to prevent PAP degradation, leading to the activation of PAP-mediated retrograde signalling and up-regulation of stress responsive genes, resulting in enhanced oxidative stress tolerance.

## Discussion

### High throughput screening and identification of a novel inhibitor for modulation of chloroplast signalling

Here we used a target-based approach to screen small molecules that exhibit increased inhibitory potency against AtSAL1 protein compared to the previous known inhibitor, Li^+^ ion (6, 46, 47). SAL belongs to a family of phosphatases that originated from a common ancestor, including fructose bisphosphatases (FBPase), inositol polyphosphate 1-phosphatase (IPPase), and inositol monophosphatase (IMPase) (48). The malachite green assay in our primary screen has been commonly used to assess inositol phosphatase activity (39), and to identify inhibitors of inositol phosphatases in drug screens (49), but its absorbance readout limits sensitivity (50). Our secondary screen accounted for this limitation by employing the highly sensitive and specific quantification of AMP and PAP via HPLC (2, 24), resulting in the identification of 97 putative compounds capable of inhibiting AtSAL1 activity *in vitro*.

A significant number of the identified inhibitors exhibit aggregation and were observed to be detergent-sensitive (Figure S3b). This includes the most potent inhibitors, such as **V77** and **V61** (Figure S3c), which contain a tryptoline ring, the most prevalent core scaffold (Group 1), found in 33% of the hit compounds. Notably, compounds with this core structure have previously been reported to aggregate (51). Colloidal aggregation is a well-recognized artifact in high-throughput screening (HTS) libraries, where aggregates can non-specifically inhibit or, in some cases, activate proteins (52). Likewise, pan-assay interference compounds (PAINS) can generate false positives through nonspecific interactions, and certain PAINS are also known to act as colloidal aggregators (53). Because of these effects, both colloidal aggregation and PAINS require careful consideration in medicinal chemistry programs. To ensure HTS hits reflect genuine interactions, several strategies to mitigate compound aggregation have been developed. These include computational identification of potential aggregators (54), using non-ionic detergents like Triton X-100 or Tween-20 to attenuate activity, detecting aggregated particles via dynamic light scattering, counter-screening against compound aggregation-sensitive enzymes, and assessing inhibition profiles for noncompetitive behaviour or attenuation with increased target protein concentration (55) Interestingly, protein aggregation plays a natural role in plant stress responses, particularly through liquid–liquid phase separation (LLPS), which underpins the formation of membraneless organelles like stress granules which can regulate stress signalling (56, 57). In the future, it may be of interest to utilise the identified aggregators herein (Figure S2) as tools to induce protein aggregation in plant cells.

We identified a sole compound, **V20**, as a genuine inhibitor of AtSAL1 activity, exhibiting 3 times greater potency than Li^+^. **V20** is a competitive inhibitor and does not share structural similarity with previously identified inositol phosphatase inhibitors (49, 58–64). Previously reported inhibitors of inositol monophosphatases are substrate analogues (competitive inhibitors) (58, 61, 62) and non-competitive inhibitors (63, 64). One class of competitive inhibitors was reported to exert their activity by chelating active site Mg^2+^ with a dihydroxytropolone (59). The discovery of **V20** introduces a novel class of compounds that inhibit the AtSAL1 phosphatase by a different mechanism (as discussed below).

### Binding sites and mechanism of action of V20

MD simulations identified two potential binding sites for **V20** on AtSAL1, designated as CBS1 and CBS2 (Figure 3; Figure S7). Chemical analogues and mutagenesis data provided further insights into the mechanism of **V20**-mediated inhibition. Based on the observed structure–activity relationships (SAR) of **V20** and its analogues and binding of **V20** in MD simulations, CBS1 is likely the primary binding site for inhibitory activity, while CBS2 may serve as an alternative binding mode that accommodates more flexible **V20** conformations. For example, substituting chlorine (Cl) to fluorine (F) in Ring A, which mainly has interactions with non-polar residues, improved inhibitory potency by 10% (Figure 2B). This enhancement is likely due to the smaller size of F and its higher electronegativity, enabling a tighter and more complementary fit within the hydrophobic binding pocket (Figure 3E iii).

The amide (NH) group in the triazole carboxamide served as a hydrogen bond donor to the backbone carbonyl of the protein and side chains of S49 and Q25 at CBS1 and CBS2, respectively (Figure 3C). This interaction appears to be critical, as evidenced by the reduced potency of the **V20-17** analogue (Figure 2B), in which substitution of the NH with oxygen eliminates N−H donor capacity and disrupts the stabilising hydrogen bonds. The replacement of the triazole ring (Ring B) with an oxadiazole in analogues such as **V20-5** and **V20-6** significantly reduced activity, likely due to the absence of the polarised C−H in the oxadiazole motif. Within the triazole scaffold, C–H bond polarisation imparts significant partial positive charge onto the H (Figure S12), and enables it to participate in H-bonding type interactions with S267 and G48 residues at CBS1 and CBS2, respectively.

Modifications of the Ring C moiety showed correlation with binding interactions at both CBS1 and CBS2. The phenyl of Ring C engages in π–π stacking with the aromatic residues F192 at CBS1 and Y148 at CBS2 (Figure 3C and D). These interactions were strengthened by electron-withdrawing substituents on the benzene ring, which is consistent with Hunter–Sanders model of π–π interactions (65). Electron withdrawing substituents reduce electron density in the π-system, thereby minimizing electrostatic repulsion and enhancing stacking potential. This also aligns with established roles of benzene ring polarizability in modulating stacking interactions (66, 67).

Although Ring C is solvent-exposed at CBS2, an independent simulation (Replicate 2, Cluster 4; Figure S6) revealed an alternative pose in which this phenyl ring is oriented inward. This conformation was maintained with 85.5% ligand residency for 0.8 µs and persisted through the end of the simulation (Figure S7C), suggesting that **V20** may adopt multiple binding modes at CBS2. Further simulations are needed to assess the stability and relevance of this alternative binding pose.

Site-directed mutagenesis identified four key residues that modulate the inhibitory potency of **V20**: K140, D258, S35, and D146. While these residues do not directly contact **V20** at either CBS1 or CBS2, structural analysis revealed that several are part of a cluster of charged residues (K140, D258, E229, E232, R256, and K262) that forms when **V20** is bound to AtSAL1. This cluster was not observed in MD simulations with PAP bound at the active site (Figure S11), suggesting that **V20** may exert its inhibitory effect by inducing conformational changes that promote stabilizing interactions among these charged residues. Interestingly, in the PAP-bound simulation, three of these charged residues (K140, D258, and K262) interact directly with PAP, leading to its binding at the active site (Figure S4B). Upon **V20** binding, the interactions of these residues with other charged residues may render them unavailable for PAP interaction, leading to inactivated conformation(s) of AtSAL1. Additionally, **V20** binding appears to influence the conformation of the active site pocket by stabilizing the positioning of Loop 1 over the catalytic site essential for PAP hydrolysis (Figure 3C; Figures S9B, S10). Disruption of these interactions may alter local positioning or flexibility of Loop 1, thereby impairing **V20**’s inhibitory effectiveness.

### V20 functions in vivo as a modulator of chloroplast retrograde signalling and oxidative stress tolerance

We found that a single application of exogenous **V20** to wild type *Arabidopsis* increased PAP levels, increased the expression of PAP-regulated oxidative stress responsive genes, and enhanced whole-plant tolerance to MV oxidative stress (Figure 4). These results indicate the efficacy of **V20** as a AtSAL1 inhibitor *in vivo* and demonstrates a proof-of-concept that novel chemical tools can be developed to selectively ‘drug’ and manipulate chloroplast signalling pathways. While HTS has been used to identify compounds involved in plant abiotic stress responses (68–70) and cellular processes like exocytosis (71), to our best knowledge, this is the first time HTS has been performed to control chloroplast-to-nucleus signalling which is a key modulator of plant environmental sensing (1, 72–74).

Previous approaches to studying SAL function have relied heavily on genetic mutants, which often exhibit strong pleiotropic effects, making it difficult to distinguish direct targets of SAL-PAP signalling from secondary effects of its knockout or knockdown. Expression of different SAL variants in *sal* null mutants has demonstrated that varying PAP levels can lead to partial or complete complementation of the mutant phenotype (2, 8, 9). However, these experiments lack control over the temporal accumulation of PAP, which limits the ability to study its dynamic role in signalling. RNA interference (RNAi) approaches failed to elevate PAP levels despite suppressed SAL protein levels (22). Attempts to use exogenous PAP feeding have also been limited in scope. While PAP-induced stomatal closure has been observed in guard cells in isolated *Arabidopsis* epidermal peels and after petiole feeding over a one-hour period (24), exogenous PAP application onto leaves did not induce transcriptional changes (18). The reliance on genetic mutants with pleiotropic effects raises important questions about the specificity of biological processes identified to interact with SAL. A chemical inhibitor specific to SAL is significant because it addresses key limitations in the current methods for studying this evolutionarily conserved pathway. A chemical tool enables the modulation of SAL activity in a temporally controlled manner, potentially across multiple plant species, thus enabling the uncoupling of developmental defects from stress-induced activation of SAL-mediated pathways to reveal its mechanism(s) of action across land plants.

Several chloroplast-to-nucleus retrograde signalling pathways involve rate-limiting proteins or enzymes for signal transduction, similar to the SAL-PAP pathway. For example, the ROS-responsive MEcPP pathway relies on redox inactivation of the hydroxymethylbutenyl diphosphate synthase (HDS) enzyme, which consumes MEcPP to regulate MEcPP levels (75); the ¹O₂-responsive EXECUTOR pathway depends on FtsH protease-mediated degradation of EXECUTOR proteins (76, 77); and the dihydroxyacetone phosphate (DHAP)-mediated pathway requires the Triose Phosphate Translocator (TPT) transporter for chloroplast export of DHAP (78) Novel chemical activators or inhibitors targeting HDS, FtsH, or TPT could enable rapid, short-term modulation of these pathways in WT plants, which could supplement the insights gained from studying genetic mutants, some of which also exhibit developmental defects.

In conclusion, this study demonstrates the successful application of high-throughput, target-based screening to identify a novel potent competitive inhibitor, **V20**, of the AtSAL1 phosphatase We uncovered two previously uncharacterized functional domains of SAL adjacent to the catalytic pocket that regulate enzyme activity by mediating conformational changes upon **V20** binding to limit substrate accessibility. This provides new mechanistic insights into AtSAL1’s regulatory architecture beyond the active site. Functional assays established that **V20** effectively modulates AtSAL1 activity *in vivo*, elevating PAP levels and activating retrograde signalling pathways.

Together, these discoveries expand our molecular understanding of SAL regulation and establish a foundation for precise chemical modulation of chloroplast-to-nucleus signalling, offering a robust platform for future investigations into retrograde communication mechanisms in plants.

## Materials and Methods

### High Throughput Chemical Library Screen for SAL1 Inhibitors

High throughput assessment of recombinant *Arabidopsis thaliana* SAL1 (AtSAL1) activity in the presence of small molecules was carried out using a colorimetric assay based on malachite green (39), which detects the production of P_i_ from AtSAL1-mediated PAP degradation. In brief, 0.2 µg of SAL1 protein was incubated in 96-well plates containing either DMSO (positive control for AtSAL1 activity with no inhibitors), 500 µM LiCl (inhibition control for AtSAL1 activity) or 500 µM small molecules from DIVERSET® library (Chembridge, USA) in 100 µL of 2X reaction buffer (500 mM Tris-MES, pH 7.5; 100 mM Mg(OAc)_2_) for 10 minutes at room temperature. An equivalent volume of PAP was added to a final concentration of 85 µM PAP, followed by incubation at room temperature for 30 minutes with shaking. The enzymatic reaction was terminated by addition of 50 µL colour reagent (0.6 g/mL malachite green in 6 N H_2_SO_4_, 3.25% ammonium molybdate, 0.17% Tween-20). After incubation at room temperature for 10 min, the resulting malachite green-phosphomolybdate complex was quantified by absorbance at 623 nm. All the pipetting, incubation and measurement steps above were performed using a JANUS^TM^ Automated Workstation (Perkin Elmer, USA) for throughput and reproducibility. Putative AtSAL1 inhibitors from the DIVERSET library were identified from wells with significantly lower A623 relative to the DMSO solvent control.

### Synthesis of AtSAL1 Lead Inhibitor, V20, and Analogues

Chemical synthesis of **V20** and analogues **V20-2** – **V20-27** was conducted by the Malins group at the Research School of Chemistry, ANU. Full experimental procedures; single-crystal X-ray analysis of **V20**; purity analyses; and ^1^H and ^13^C NMR spectra of **V20** and analogues **V20-2** – **V20-27** are reported in the Supporting Information (SI).

### Small Molecule Single-Crystal X-ray Diffraction (CX)

Single-crystal X-ray diffraction data for **V20** were collected at 150 K with Mo Kα radiation (λ=1.54184 Å). The crystal belonged to the triclinic space group P −1, with unit cell dimensions (a=5.4391 Å, b=10.4877 Å, c=16.3127 Å) and angles (α=72.894 °, β=86.189 °, γ=81.635 °). Data completeness was 99.3% to a maximum θ of 66.57°. Absorption corrections were applied using a multi-scan method. Final refinement statistics included *R_1_*=0.0530 for observed data and w*R_1_*=0.1106, with a goodness of fit of 0.991. Molecular formula was determined as C19 H17 Cl N4 O3, with a calculated density of 1.453 g/cm³ and absorption coefficient μ=2.174 mm^−1^. The dataset included 3092 reflections and 249 parameters were refined. The CIF and CheckCIF validation reports are provided in the SI. The structure has been deposited in the Cambridge Structural Database (CSD; deposition code: 2488420).

### Mutagenesis of the AtSAL1 cDNA in Plasmids

Mutagenesis was performed using primers containing point mutations (Table S2) to change the residue of interest into an alanine, with the QuikChange II XL Site-Directed Mutagenesis Kit (Stratagene, USA) as per the manufacturer’s instructions. The template plasmid DNA used in the PCR reaction was the pHUE vector containing the mature full-length WT AtSAL1 cDNA without its transit peptide. All point mutations were confirmed by Sanger sequencing at the Biomolecular Resource Facility (BRF), Australian National University.

### Recombinant Protein Purification from E. coli

The full-length sequence of AtSAL1, excluding the chloroplast transit peptide sequence, was expressed in *Escherichia coli* BL21 (DE3) cells as described previously (14). Briefly, the protein was expressed as an N-terminal polyhistidine-tagged ubiquitin fusion protein using the pHUE expression vector. Expression was induced with isopropyl β-D-1-thiogalactopyranoside (IPTG). The fusion protein was purified from the soluble cell lysate using Ni-NTA His-Bind Resin (Novagen) via immobilized metal affinity chromatography according to the manufacturer’s protocol. The polyhistidine-ubiquitin tag was subsequently cleaved by digestion with the deubiquitinylating enzyme Usp2. The cleaved tag was removed by passing the protein solution back over the Ni-NTA resin. The purified, untagged AtSAL1 protein was concentrated and buffer-exchanged into a storage buffer (50 mM Tris-HCl, pH 8.0, 150 mM NaCl, 20 mM KCl, 1 mM MgCl₂, and 15% (v/v) glycerol) and stored at −80°C until further use. Protein purity was assessed to be >95% by SDS-PAGE. For crystallisation experiments, before being buffer exchanged, the protein was further purified by size exclusion chromatography using a HiLoad 26/60 Superdex-200 column (GE Healthcare).

### Assay of Recombinant SAL1 Protein Activity

Enzyme kinetics and AtSAL1 activity upon inhibitor treatment against PAP were assayed by pre-incubating 0.2 µg of AtSAL1 protein with each inhibitor (concentration specified in results) for 10 minutes at room temperature in 2X reaction buffer (500 mM Tris-MES, pH 7.5; 100 mM Mg(OAc)_2_). An equivalent volume of PAP was added to a final concentration of 85 µM PAP (unless otherwise specified). The reaction mixture was incubated at room temperature for either 30 minutes for the screening of inhibitory activity by small molecules or 5 minutes for IC_50_ assays and determination of Michaelis-Menten kinetics, with varying PAP and inhibitor concentrations. The reaction was stopped by flash-freezing in liquid nitrogen, and the AMP produced by SAL1 degradation of PAP was quantified using High Performance Liquid Chromatography (HPLC). The enzymatic activity of SAL1 was calculated relative to the DMSO control (no inhibitor). The IC50 values, Michaelis-Menten kinetics, and Lineweaver-Burk parameters were determined using GraphPad Prism software. In detergent sensitivity assays, Triton X-100 was added to the reaction after pre-incubation with the inhibitor to a final concentration of 0.01% and 0.1% (*v*/*v*).

### Quantification of Adenosine in vitro

Quantification of PAP and other adenosine compounds was performed as described previously by (2). Briefly, 100 µL of the enzymatic reaction solution was mixed with 810 µL CP buffer (620 mM citric acid monohydrate; 760 mM disodium hydrogen phosphate dihydrate; pH 4.0) to denature the AtSAL1 protein. The adenosines were then derivatised with 80 µL of chloroacetaldehyde (CAA) for fluorescent detection of compounds. CAA selectively reacts with the purine ring of adenosine nucleotides, forming stable adducts called ethenoadenosine derivatives, which fluoresce upon excitation at 280 nm and emit at 410 nm (79). 5 µL of the derivatised compounds were then injected into an Agilent LC1100 High-Performance Liquid Chromatography (HPLC) machine, fractionated on a Kinetex butylated C-18 column (Phenomenex), and detected fluorometrically. The run for each sample took around 16 minutes, and the solvent gradient to separate adenosine compounds was as follows: equilibration of the column for 0.05 minutes with 95% (*v*/*v*) of buffer A (5.7 mM tetramethylammonium hydrogen sulfate; 30.5 mM potassium dihydrogen phosphate; pH 5.8) and 5% (*v*/*v*) buffer B (67% (*v*/*v*) acetonitrile and 33% (*v*/*v*) buffer A), linear gradient for 13.45 minutes up to 50% (*v*/*v*) of buffer B, and re-equilibration for 1.8 minutes with 5% (*v*/*v*) buffer B. Chromatograms of detected fluorescence and column pressure over time were recorded for each run and analysed with the Agilent LC1100 ChemStation software for quantification of peak areas. Commercial standards of derivatised AMP and PAP (Sigma-Aldrich 01930 and A5763, respectively) were also run on the HPLC during each experiment (Figure S1E, F) to construct calibration curves for quantifying the amount of AMP and PAP in each sample.

### Protein Crystallisation

Crystals of the purified AtSAL1 were grown at room temperature using the hanging-drop vapor diffusion method. The protein solution was concentrated to approximately 20 mg/mL. The reservoir solution consisted of 28% (w/v) PEG 3350, 0.15 M ammonium sulfate, and 0.1 M HEPES buffer at pH 8.2. Drops were prepared by mixing equal volumes (1 µL) of the protein and reservoir solutions. Crystals appeared within 3-5 days.

### Diffraction Data Collection and Structure Refinement

Diffraction data were collected at the Australian Synchrotron MX2 beamline (80). Crystals diffracted to a resolution of 2.60 Å and belonged to the hexagonal space group *P*6₁22, with unit cell dimensions of a=142.97 Å, b=142.97 Å, c=75.14 Å (Table S1). Data were integrated with XDS (81) and scaled with AIMLESS (82) as implemented in the CCP4 suite. Two datasets, collected from different regions of the crystal were merged in AIMLESS, resulting in high multiplicity and *R*_merge_, given the tendency for R_merge_ to increase with increasing multiplicity (83). The CC1/2, I/σI, and R_pim_ values were acceptable in the outer shell (84). Unlike the previously reported structure, these crystals were not affected by merohedral twinning. The structure was solved by molecular replacement using PHASER (85). Refinement was carried out using phenix.refine (86), resulting in final *R*_work_ and *R*_free_ values of 0.208 and 0.248, respectively. The final model, deposited in the RCSB PDB under accession code 8F9Y, shows significant improvement in the resolution of surface loops and side-chain conformations over previous low-resolution model.

### Plant Materials and Growth Conditions

*Arabidopsis thaliana* WT and *atsal1* mutant plants used in this study are in the ecotype Col-0 background. The *sal* null mutants include *altered expression of APX2 8* (*alx8*), which has a single point mutation (G217D) in the *SAL* (At5G63980) gene (2, 3). The plants were grown in soil in individual pots at 23 °C with a photoperiod of 16 hours light and 8 hours dark.

### Determination of Oxidative Stress Tolerance of Arabidopsis Leaf Discs Induced by Methyl Viologen Treatment

25-day-old *Arabidopsis* plant leaves were punched into discs and put into 100 µL of buffer (1 mM PIPES (pH 6.25), 1 mM KCl, 1 mM sodium citrate, 1 mM sucrose, and 0.1% Tween, with or without 250 µM inhibitor) in a 96-well plate. The tissues were vacuum infiltrated for 10 minutes, and methyl viologen (MV) or buffer solution (untreated control) was added to a final concentration of 0.5 µM and incubated overnight in the dark. Fv/Fm was measured for up to 10 hours using the Imaging-PAM chlorophyll fluorometer (Walz). Chlorophyll fluorescence measuring protocols involved repetitive 1-hour periods of blue actinic light (450 nm, 66 µmol m^−2^ s^−1^), followed by 20 minutes of dark adaptation (87). During each 1-hour period, chlorophyll fluorescence under light was recorded every 2 minutes. After each 1-hour actinic light period, a 20-minute dark adaptation was followed, during which F_o_ and F_m_ were measured, and the next light cycle was initiated.

### Preparation of V20-Containing Liposomes for Treatment on Whole Plants and Protoplasts

**V20** (50 μL, 20 mM, in 100 % ethanol) was mixed with lipids (200 μL of 125 μg/μL L-α-phosphatidylcholine (Sigma Merck 840058P) in 100 % ethanol, and 50 μL of 20 mM DEPC-PE2000 (Sigma Merck 880120P) in 100 % ethanol). For the mock treatment, **V20**-ethanol was substituted by 50 μL DMSO. After incubation with shaking (42 °C, 400 rpm) for approximately 5 minutes, the mixture was concentrated by vacuum centrifugation at 45 °C until ∼50 μL remained. 1000 μL of 0.5x phosphate-buffered saline (PBS) was then added to each tube to facilitate liposome formation. Samples were mixed by flicking and vortexing, and then sonicated on ice (30 % amplitude, 2-second pulse, 5 minutes per round for 3 rounds). Samples were placed on ice for 2 minutes between each round of sonication. The liposomes were then further diluted with 1,300 μL of 0.5x PBS, resulting in a final volume of 2.3 mL. The liposomes were stored at 4 °C for no more than a week. The liposome solutions were warmed to room temperature and diluted with PBS to a final volume of 4 mL (final concentration of **V20**: 250 μM) prior to application onto plants.

### Protoplast Isolation and Treatment with V20 and PAP

Thin (∼1 mm) strips of 3-4-week-old *Arabidopsis thaliana* leaves were prepared in digestion buffer (DB) [20 mM MES-KOH (pH 5.7), 600 mM sorbitol, 10 mM KCl, 1.2 % (*w*/*v*) cellulase RS (Yakult), 0.6 % (*w*/*v*) macerozyme R-10 (Yakult), 10 mM CaCl_2_, 10 mM dithiothreitol (DTT), 0.1 % (*w*/*v*) bovine serum albumin (BSA)] and vacuum infiltrated for 15 mins. The leaf strips were incubated in the dark at ∼25 °C and 75 rpm for 1.5 hours. Protoplasts were harvested by adding cold wash buffer (WB) [20 mM MES-KOH (pH 5.7), 600 mM sorbitol, 10 mM KCl, 10 mM CaCl_2_, 10 mM DTT, 0.1 % (*w*/*v*) BSA] to the digestion mixture, and gently swirled. The mixture was then transferred onto a 100 μm mesh using a 1000 µL cut micropipette tip and filtered to remove undigested leaf material. The protoplast yield was enhanced by repeating the filtration with additional WB on the unfiltered leaf debris. Mesophyll protoplasts in the filtrate were pelleted by centrifugation for 5 minutes at 300 g and 4 °C in a swing bucket rotor. The crude pellet of mesophyll protoplasts was purified by resuspending in 5 mL of Dex15 [20 mM MES-KOH (pH 5.7), 600 mM sucrose, 10 mM KCl, 10 mM CaCl_2_, 10 mM DTT, 15 % (w/v) Dextran T40], which was then overlaid with 3 mL of Dex10 [same composition as Dex15 but with 10 % (w/v) Dextran T40 instead of 15 %], and 3 mL of cold WB. The gradient was centrifuged for 10 minutes at 300 g and 4 °C in a swing bucket rotor. Mesophyll protoplasts were retrieved from the WB-Dex10-Dex15 interface, washed with WB and pelleted (5 minutes at 300 g and 4 °C in a swing bucket rotor). The purified protoplasts were resuspended in 1-2 mL WB. Cell count and viability was determined using a haemocytometer and a Nikon Eclipse 50i light microscope (Exposure: 0.11, Gain: 6, Gamma: −16, Saturation: 5, Magnification: 10x).

Approximately 150,000 protoplasts per replicate were treated with either DMSO alone, 100 µM PAP alone, 250 µM **V20** alone, or 100 µM PAP and 250 µM **V20** in WB for 16 hours. Protoplasts were collected via centrifugation, flash-frozen and their PAP content determined as described previously.

### Gene Expression in Response to V20 Treatment and Methyl Viologen-Induced Oxidative Stress

18-21 days old *Arabidopsis* plants were sprayed with 330 μL per plant of either 250 μM AtSAL1 inhibitor **V20** in liposomes or mock solution (liposomes alone) with an air brush (Iwata). After overnight incubation in the growth chamber, the plants were then treated with either 100 μM MV + 0.1% Tween-20 or 0.1% Tween-20 alone, incubated in the dark for 2 hours, and then exposed to growth light for 2.5 hours. Maximal photosystem II efficiency (Fv/Fm) was measured at 0, 30, 60, 90, 120, and 150 minutes post exposure to light, prior to tissue harvest for gene expression analysis.

Gene expression in inhibitor- and MV-treated plants was analysed using quantitative RT-PCR. Total RNA was extracted from tissue samples using the RNeasy Plant Mini Kit (Qiagen) and treated with Invitrogen™ DNase I (ThermoFisher Scientific) to remove genomic DNA. For the RT-PCR reaction, 100 ng of RNA per biological sample was used. RT-qPCR was carried out with Power SYBR™ Green RNA-to-CT™ on a QuantStudio™ 6 Flex Real-Time PCR System (Applied Biosystems™). Cycling conditions are as per manufacturer recommendations, consisting of a reverse transcription step at 48 °C for 30 minutes, activation of AmpliTaq Gold® DNA Polymerase at 95 °C for 10 minutes, 40 amplification cycles of denaturing at 95 °C for 15 seconds and annealing/extension at 60 °C for 1 minute. A melt curve analysis was performed at the end to assess amplicon specificity. Expression of *PROTEIN PHOSPHATASE 2A SUBUNIT A3 (PP2AA3)* gene was used as the internal reference for normalisation. RT-PCR primer sequences are provided in Table S3.

### Molecular Dynamics Simulation

The molecular dynamics simulation systems were built using the AtSAL1 protein (PDB ID: 8F9Y) with four molecules of PAP, V20, V30, or V01, utilizing VMD software (version 1.9.4). Systems were solvated and ionised with water molecules and 150 mM K⁺ and Cl⁻ ions to neutralize net charge. Parameters for small molecules, protein, water, and ions were assigned via the CHARMM36 general force field (CGenFF version 2.2.0) with penalty scores checked to ensure reliable force field description. Small molecule ligands were modelled with MarvinSketch software (version 22.11.1).

CHARMM parameters were converted to AMBER-compatible formats using the CHAMBER tool, enabling accelerated molecular dynamics simulations in AMBER and AmberTools (AMBER20). Hydrogen mass repartitioning (HMR) was performed using ParmEd (version 3.4.0) to redistribute hydrogen mass, allowing an increased time step (to 4 femtoseconds) without stability loss. Trajectory analyses were conducted using VMD software for visualisation and molecular dynamic assessments. Further details on simulation protocols, analysis methods, and parameterisation can be found in the SI.

### Statistical Analysis

Analysis of variance (ANOVA) was used to assess significant differences (p<0.05) between three or more sample groups, and one-way ANOVA was used when only one parameter (such as genotype) was varied. Multiple-way ANOVA was employed when two or more parameters were changed. The two-sample Student’s t-test was utilised when comparing two sample groups. GraphPad Prism 7.0 (GraphPad Software, USA) or R3.08 (www.r-project.org) was used for all statistical analyses.

## Supporting information

All Supplemental Data

## Acknowledgments

This work was funded by the Australian Research Council (ARC) Laureate Fellowship (FL190100056 to BJP); ARC Discovery Early Career Researcher Award (DE210100466 to KXC); ARC Future Fellowship (FT240100010 to LRM); ARC Centre of Excellence (CoE) for Innovations in Peptide and Protein Science (CE200100012 to BDS, LRM, LLT, CJ), ARC CoE in Synthetic Biology (CE200100029 to CJ); ANU-UC Connect Ventures Discovery Translational Funds 2016 and 2017 to BJP and KXC; Australian Government Department of Education Endeavour Leadership Program (6975_2018) and CSIRO Synthetic Biology Future Science Platform (MC); ANU HDR Scholarship (NCK); ANU Honours Scholarship (SY). Diffraction data for crystal structure was collected using the MX2 beamline at the Australian Synchrotron, part of ANSTO, and made use of the Australian Cancer Research Foundation (ACRF) detector. This research was undertaken with the assistance of resources from the National Computational Infrastructure (NCI Australia), an NCRIS enabled capability supported by the Australian Government. We thank Ran Ashkenazi, Zhong-Yu Wang and Rebecca Frkic for technical assistance with the V20 application, HTS experiments and crystallisation work, respectively.

