## Supplementary material for "High-throughput screening and structure-guided design of small molecules enable modulation of SAL-PAP stress signaling": All Supplemental Data

**This PDF file includes:**

Supporting text
Figures S1 to S12
Tables S1 to S2
SI References

### Supporting Information Text

### Supplementary Methods

*Molecular Dynamics Simulation*

**Preparation of simulation systems.** Simulation systems were constructed using the AtSAL1 protein (PDB ID: 8F9Y) and four molecules of either PAP, **V20**, **V30**, or **V01**, using VMD software (version 1.9.4). Each system was solvated and ionised with water molecules and ions (150 mM $K^+$ and $Cl^-$) to neutralise the system net charge. In each system 4 copies of the small molecule were included, randomly located and oriented in the aqueous regions at least 20 Å away from the AtSAL1 protein.

Parameters for the small molecules, protein, water, and ions (including the $Mg^{2+}$ cofactor) were defined using the CHARMM36 general force field (version 2.2.0) (1), which includes atomic charges, van der Waals interaction parameters, and covalent bond descriptions (including dihedral angles). Initial 3D coordinates of the small molecule ligands were generated using MarvinSketch software (version 22.11.1). These molecules were then parameterised using the CHARMM General Force Field (CGenFF version 2.2.0) and manually inspected to ensure the absence of high-penalty parameters.

Li–Merz OPC-optimised parameters were used for the $Mg^{2+}$ ion (2), while parameters for $K^+$ and $Cl^-$ were taken from Joung and Cheatham (3). Water molecules were represented using the TIP3P (Transferable Intermolecular Potential with 3 Points) model, which treats water as a rigid, non-vibrating molecule with fixed OH bond lengths and HOH bond angles (4).

CHARMM force field parameters were converted to AMBER-compatible formats using the CHAMBER tool (5) to enable accelerated molecular dynamics simulations with the AmberTools package and AMBER molecular dynamics software (AMBER20). Hydrogen mass repartitioning (HMR) was applied to redistribute some mass from heavy atoms to bonded hydrogens, allowing longer simulation time steps without compromising stability (6). HMR was performed on the topology files using the ParmEd package (version 3.4.0) (7).

**Accelerated Molecular Dynamics Simulations.** The system was then minimised to a lower energy state, gradually heated from 0 K to 310 K to prevent excessive and sudden solute fluctuations while restraining the protein backbone atoms and alpha carbons of ligands with a force constant of 5 kcal $mol^{-1}$ $Å^{-2}$. The restraints were then gradually relaxed over 28 ns. A short 20 ns production run of conventional molecular dynamics (cMD) was then performed to calculate the average total potential and dihedral energies, which were used to define the energy boost parameters for accelerated molecular dynamics (aMD) simulations. These boost parameters for both the total potential and torsional terms were determined based on the calculated energies and the number of atoms and residues in the system (8). aMD simulations were run on the Gadi supercomputer (National Computational Infrastructure (NCI Australia)). After the system was equilibrated, it was simulated using AMBER20 (9) with a time step of 4 femtoseconds (compared to 2 femtoseconds in cMD) at a temperature of 310 K maintained by a Langevin thermostat, and a pressure of 1 atm maintained with a Langevin piston. Particle-Mesh Ewald (PME) method was used to calculate long-range electrostatic interactions in the simulation. A cut-off distance of 12.0 Å was used for the calculation of van der Waals interactions. Periodic boundary conditions (PBCs) were applied (approximately 110 x 110 x 110 cubic/rectangular shaped box - varies slightly between systems of different ligands) to better approximate the behaviour of a system in bulk, which is more representative of macroscopic conditions.

**Analysis of Simulation Trajectories.** The simulation trajectory files were visualised and analysed using VMD software (developed by the Theoretical and Computational Biophysics Group at the Beckman Institute for Advanced Science and Technology at the University of Illinois at Urbana-Champaign) (10) for each system. Specifically, cluster analysis of ligands, plots of bond distances between the atoms of interest, and interaction energies between protein residues and the ligand were calculated using an external plugin for cluster analysis, developed by Luis

Gracia at Weill Cornell Medical College (https://github.com/luisico/clustering), and built-in NAMD Energy and bond analysis plugins. The trajectories were stripped of ions and water molecules to compress file size and to facilitate downstream analysis using CPPTRAJ (AmberTools) (11). To analyse how ligands cluster at various sites of the protein throughout the simulation, trajectories were loaded into VMD, containing one ligand at a time to observe ligand-ligand interactions and determine binding pocket occupancy. Protein backbone alignment was performed using the built-in RMSD trajectory tool before cluster analysis, with settings for clustering ligand poses within a 5 Å cut-off, resulting in a candidate binding site for the ligand. Cluster analysis outputs the trajectory frames of the top five ligand clusters. To investigate key interacting partners of the ligand at the identified binding sites, non-bond interaction energy (van der Waals and electrostatic interactions) between the amino acid residues at the binding pocket and the ligand were calculated for selected trajectory frames. The average interaction energies of various protein residues and the ligand were then plotted using GraphPad Prism software version 08.0.0 for Windows, GraphPad Software, Boston, Massachusetts USA, www.graphpad.com.

### Supplementary Figures

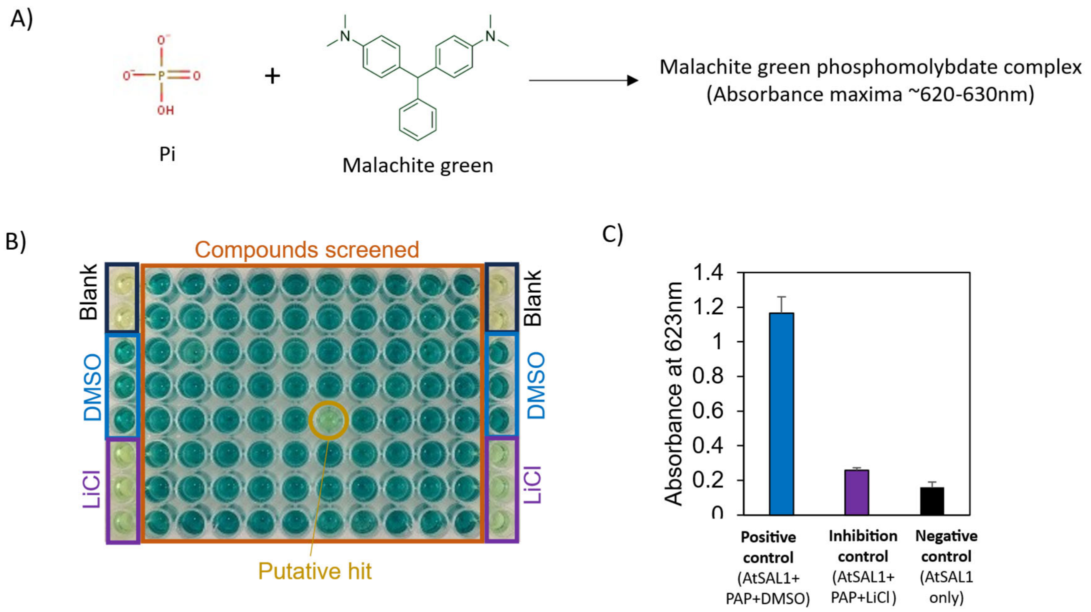


D)

| | Mean $Abs_{623nm}$ | SD | 3SD | Mean + 3SD | Mean -3SD |
|---|---|---|---|---|---|
| DMSO | 1.1663 | 0.0947 | 0.284 | 1.261 | 0.8823 |
| LiCl | 0.2575 | 0.0147 | 0.0442 | 0.3017 | 0.2428 |

| Dynamic Range, R<br>$Mean_{DMSO}$- $Mean_{LiCl}$ | Separation Band, S<br>($Mean_{DMSO}$- $3SD_{DMSO}$) – ($Mean_{LiCl}$+$3SD_{LiCl}$) | Z'-score<br>S/R |
|---|---|---|
| 0.908 | 0.580 | 0.638 |

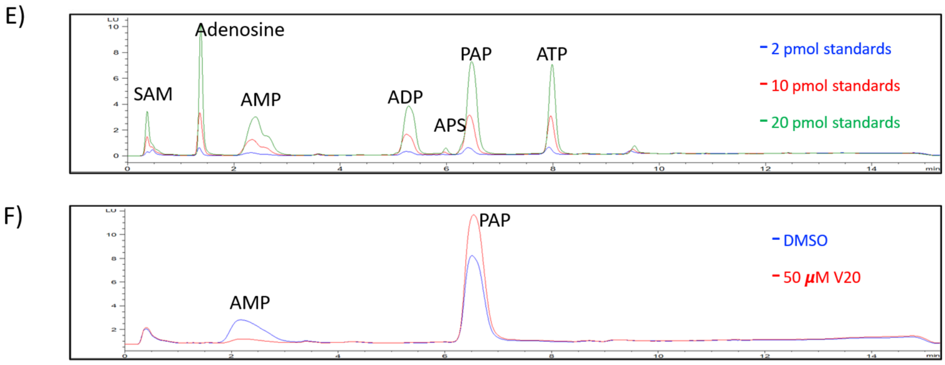


**Fig S1. High-throughput primary screening assay for AtSAL1 phosphatase activity using malachite green.**
**(A)** Schematic representation of the malachite green assay: inorganic phosphate (Pi) released from PAP hydrolysis reacts with malachite green and molybdate to form a green phosphomolybdate complex, detectable by absorbance at 620–630 nm (measured at 623 nm).
**(B)** Layout of the 96-well plate used for primary screening of a chemical library against AtSAL1 phosphatase activity. Control wells include: blanks (AtSAL1 without PAP), DMSO-treated (AtSAL1 with PAP, no inhibitor), and LiCl-treated (inhibition control, AtSAL1 with PAP and 0.5 mM

LiCl). Test compounds were added to central wells (orange box), and putative inhibitory hits were identified by both colour change and reduced absorbance (example highlighted with an orange circle).
**(C)** Absorbance values at 623 nm for control conditions (mean ± SD, *n* = 3). LiCl-treated wells show significantly reduced absorbance compared to DMSO controls, indicating inhibition of AtSAL1 activity.
**(D)** Assay performance statistics. Z'-score (0.638) was calculated using the dynamic range (R = 0.908) and separation band (S = 0.580), indicating a robust assay suitable for high-throughput screening.
**(E)** HPLC chromatograms of commercial adenosine-based nucleotide standards used to identify retention times and establish calibration curves for quantification in the secondary screening and subsequent activity assays. Standards were derivatised and analysed at concentrations of 1.34 µM (2 pmol per injection, blue), 6.7 µM (10 pmol, red), and 13.4 µM (20 pmol, green). Peaks corresponding to SAM, AMP, ADP, APS, PAP, ATP, and adenosine were annotated based on retention time.
**(F)** HPLC chromatograms showing product profiles from *in vitro* AtSAL1 phosphatase assays using 85 µM PAP as substrate. The blue trace represents the DMSO-treated control (vehicle), while the red trace corresponds to the reaction in the presence of 50 µM compound **V20**. AtSAL1 activity was inferred from AMP production (identified by retention time), with reduced AMP signal in the **V20**-treated condition suggesting inhibition of SAL-mediated PAP hydrolysis.

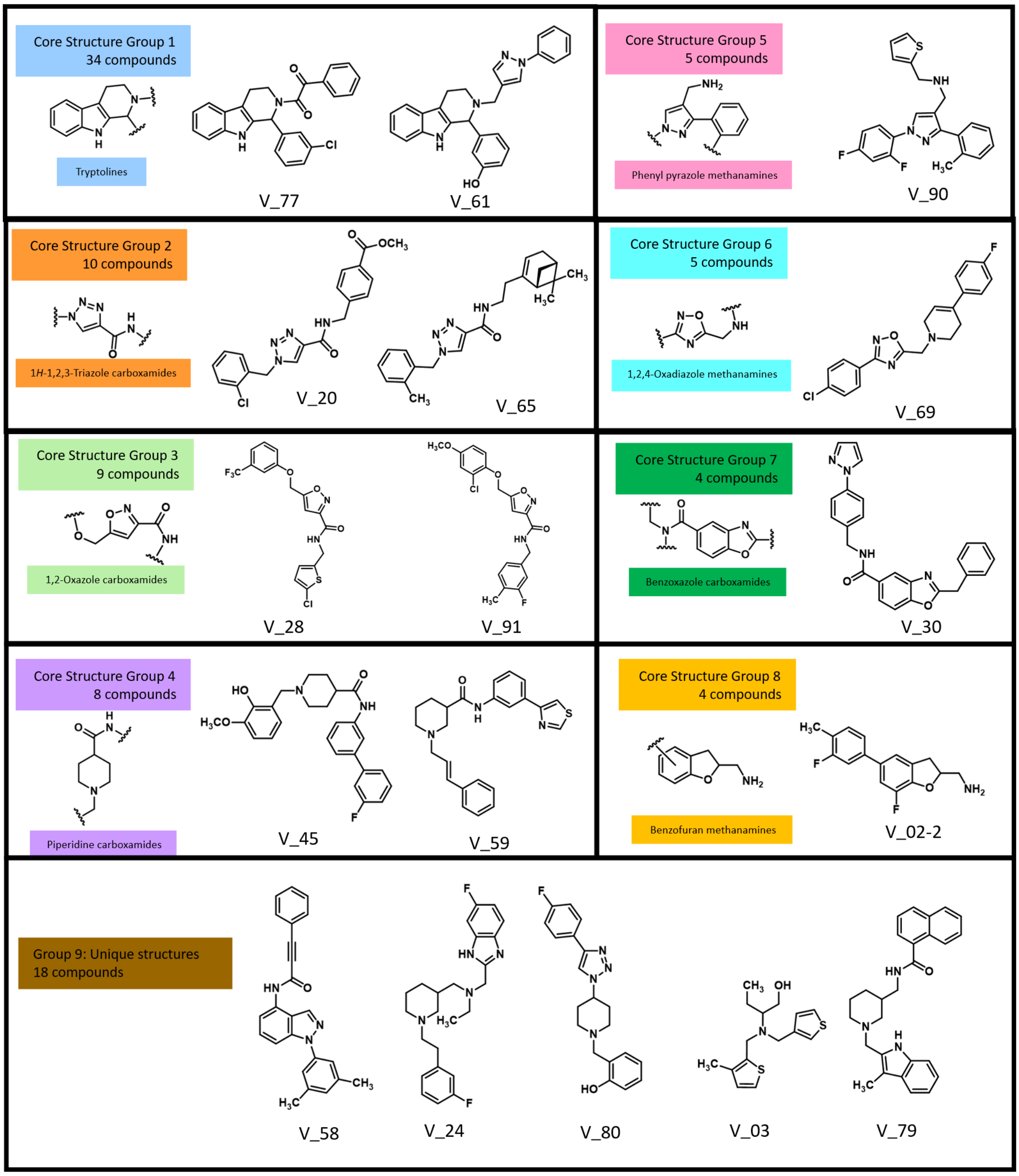


**Fig S2. Chemical diversity of AtSAL1 inhibitor candidates.**

Core scaffold diversity of 97 putative AtSAL1 inhibitors identified from secondary screening. The number of compounds sharing a particular core scaffold is indicated next to each structure. Chemical substituent positions are denoted by wavy lines. Group 9 comprises 18 molecules with unique chemical features or scaffolds shared by fewer than four compounds.

A)

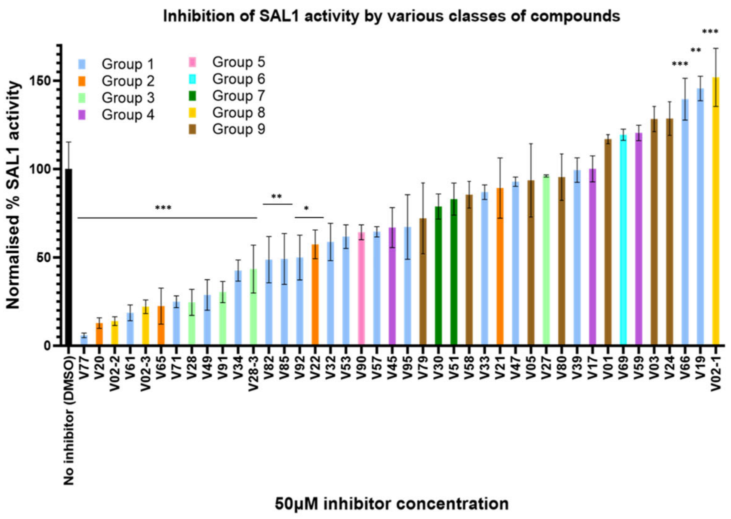


B)

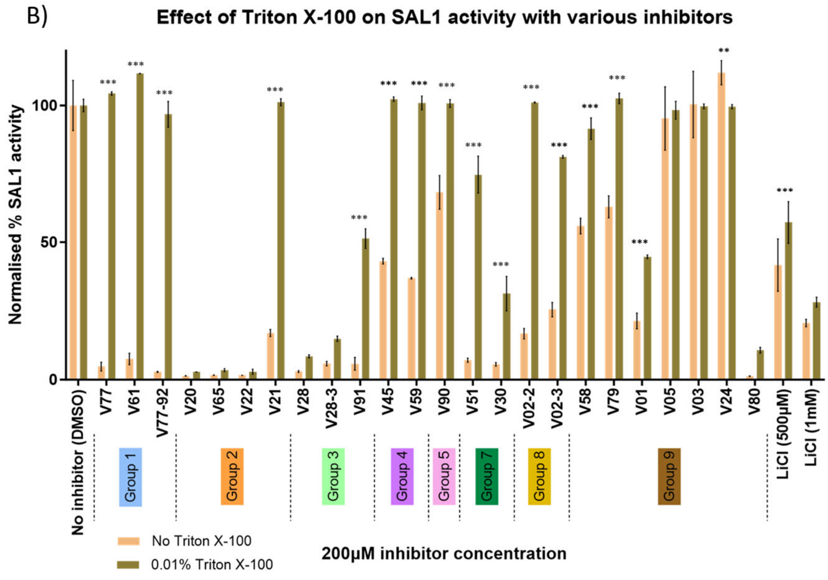


**Fig S3. Validation of AtSAL1 inhibitor candidates.**
**(A)** A total of 42 putative candidate AtSAL1 inhibitors were selected from 97 hit compounds and tested for potency at a concentration of 50µM inhibitor. The original V02 compound was not available in the Chembridge (Hit2Lead) database for testing, therefore, three structurally similar analogues (**V02_1**, **V02_2**, **V02_3**) were tested instead. AtSAL1 activity was normalised to the DMSO control, and the inhibitors were arranged in descending order of potency on the x-axis (n=3). The error bars represent standard deviations. Tukey's multiple comparison test was performed between the control and each inhibitor treatment, and the degree of significance between these two conditions is represented with asterisks (** for P ≤ 0.01, *** for P ≤ 0.001)
**(B)** AtSAL1 inhibitory activity of selected compounds tested at 200 µM in the presence or absence of 0.01% Triton X-100 (TTX-100). LiCl (0.5–1 mM) was used to assess the ionic strength tolerance of the assay. Inhibition of AtSAL1 activity was quantified by measuring AMP production and normalised to DMSO control for each TTX-100 condition (n=3). Compounds are grouped along the x-axis by scaffold class, indicated by coloured boxes. Error bars represent standard errors of the mean. Asterisks indicate statistically significant differences from the respective DMSO controls (***P ≤ 0.001, **P ≤ 0.01, *P ≤ 0.05).

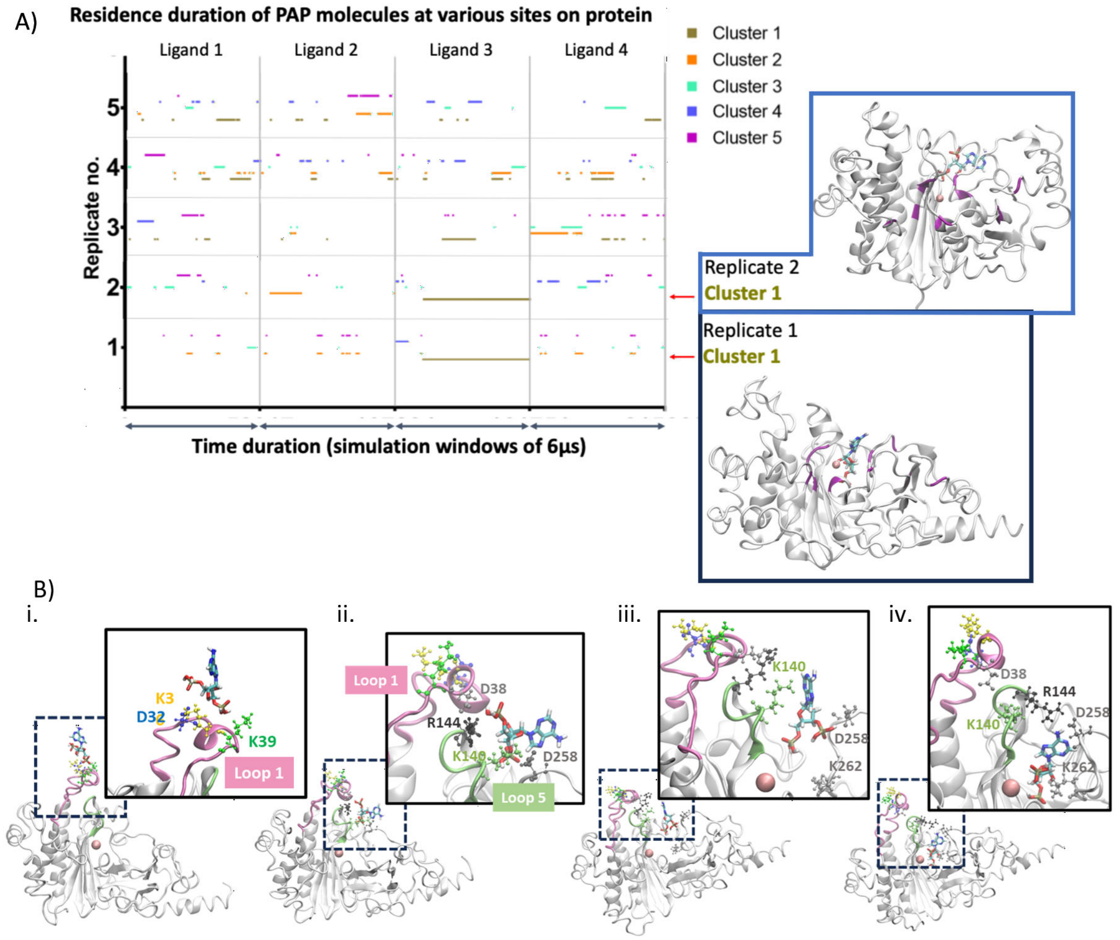


**Fig S4. Molecular dynamics simulations reveal PAP binding clusters and a dynamic substrate-guided binding mechanism in AtSAL1.**

**(A) Cluster analysis of PAP binding sites.** Molecular dynamics simulations identified recurrent PAP binding positions in AtSAL1. Clustering was based on trajectory frames where PAP conformations showed ≤5 Å RMSD. The top five clusters are shown across five independent replicates, with each simulation including four PAP ligands (x-axis). Cluster 1 (yellow) represents the most populated and stably maintained binding conformation. Replicates are separated by black lines, and clusters of interest are highlighted with red arrows. Representative protein structures with stably bound PAP molecules are shown, with PAP atoms coloured by element: nitrogen (blue), oxygen (red), phosphorus (khaki), carbon (cyan), and hydrogen (white). The predicted catalytic site of AtSAL1 is highlighted in purple.

**(B) Stepwise interaction of PAP with AtSAL1 through loop-mediated substrate guidance.** Snapshots from replicate 1 illustrate the dynamic progression of PAP binding mediated by loop 1 (pink) and loop 5 (green). The protein backbone is shown in grey. Key residues are shown in ball-and-stick format: positively charged (blue) and negatively charged (red) side chains. The following sequence was observed:

i. **Initial encounter at loop 1:** PAP phosphate groups and adenine ring interact with positively charged residues K36 and K39 on loop 1, facilitated by partial unwinding of the loop's α-helix enabled by an intraloop D32-K36 interaction.

ii. **PAP transfer to loop 5:** Loop 1 bends toward loop 5 via D38 (loop 1) and R144 (loop 5) interaction. PAP is transferred through its 3′ phosphate and K140, and later its adenine ring, anchoring it onto loop 5.

iii. **Catalytic site engagement:** Initially, the entrance to the catalytic site is occluded by an interaction between K140 and D258. PAP inserts between these residues, disrupting their interaction. The adenine ring and 3′ phosphate group engage K140 and D258, respectively, followed by interaction with K262 near the catalytic site.

iv. **Stabilisation and closure:** K140 (loop 5) subsequently interacts with D38 (loop 1), sustaining loop 1's bent conformation toward the active site. R144, which previously interacted with D38, transitions to interact with D258, contributing to catalytic site closure.

These conformational changes support a model where flexible loops coordinate substrate recognition, guide PAP entry to the active site, and stabilise PAP for catalysis.

A)

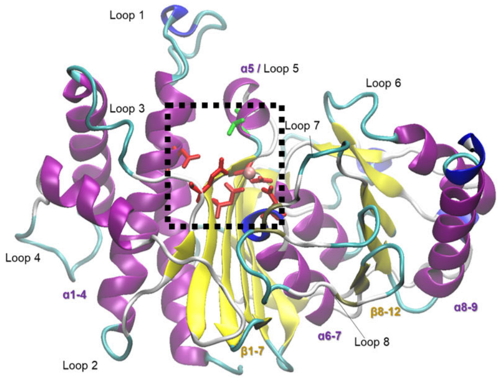


B)

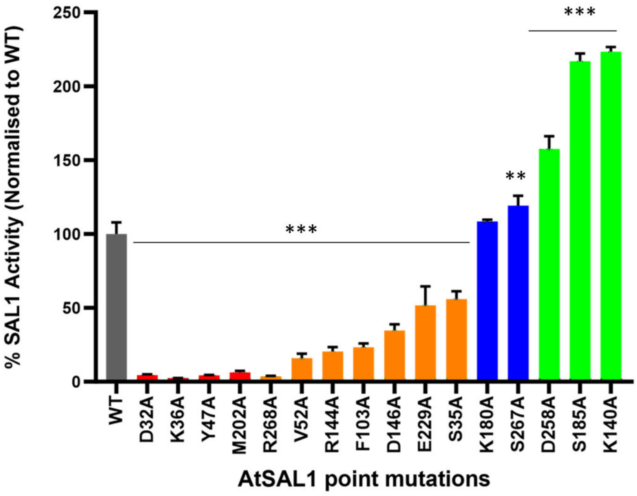


C)

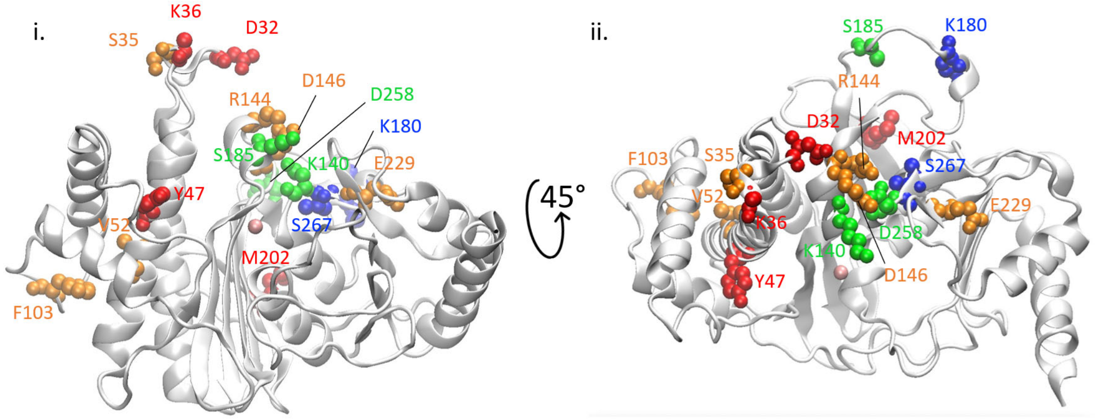


**Fig S5. Structural context of key catalytic residues in AtSAL1.**
**(A) Secondary structure and active site composition of AtSAL1.**
The crystal structure of AtSAL1 (PDB ID: 8F9Y) is shown in cartoon representation: α-helices (purple), β-strands (yellow), loops (grey). Key loops annotated. The boxed region highlights the catalytic site located at the interface of loop 1 and loop 5. A $Mg^{2+}$ ion at the catalytic site is depicted as a pink sphere. Residues involved in PAP catalysis shown in licorice: negatively charged (E71, D134, D137, D288, D46) in red; polar T139 in green.
**(B) Relative enzymatic activity of AtSAL1 single-point mutants.**

Bar graph showing the enzymatic activity of recombinant AtSAL1 point mutants, normalised to wild-type (WT) activity (n=3). Asterisks denote significant differences compared to WT enzyme activity (*P < 0.05, **P < 0.01, ***P < 0.001; Student's t-test)

**(C) Spatial distribution of mutant residues on the AtSAL1 structure.**

Mutated residues are mapped onto the AtSAL1 structure in (i) side-on and (ii) top-down views. Residues are colour-coded by mutation impact on enzyme activity: red (catalytically inactive); orange (reduced activity); blue (WT-like); green (enhanced activity).

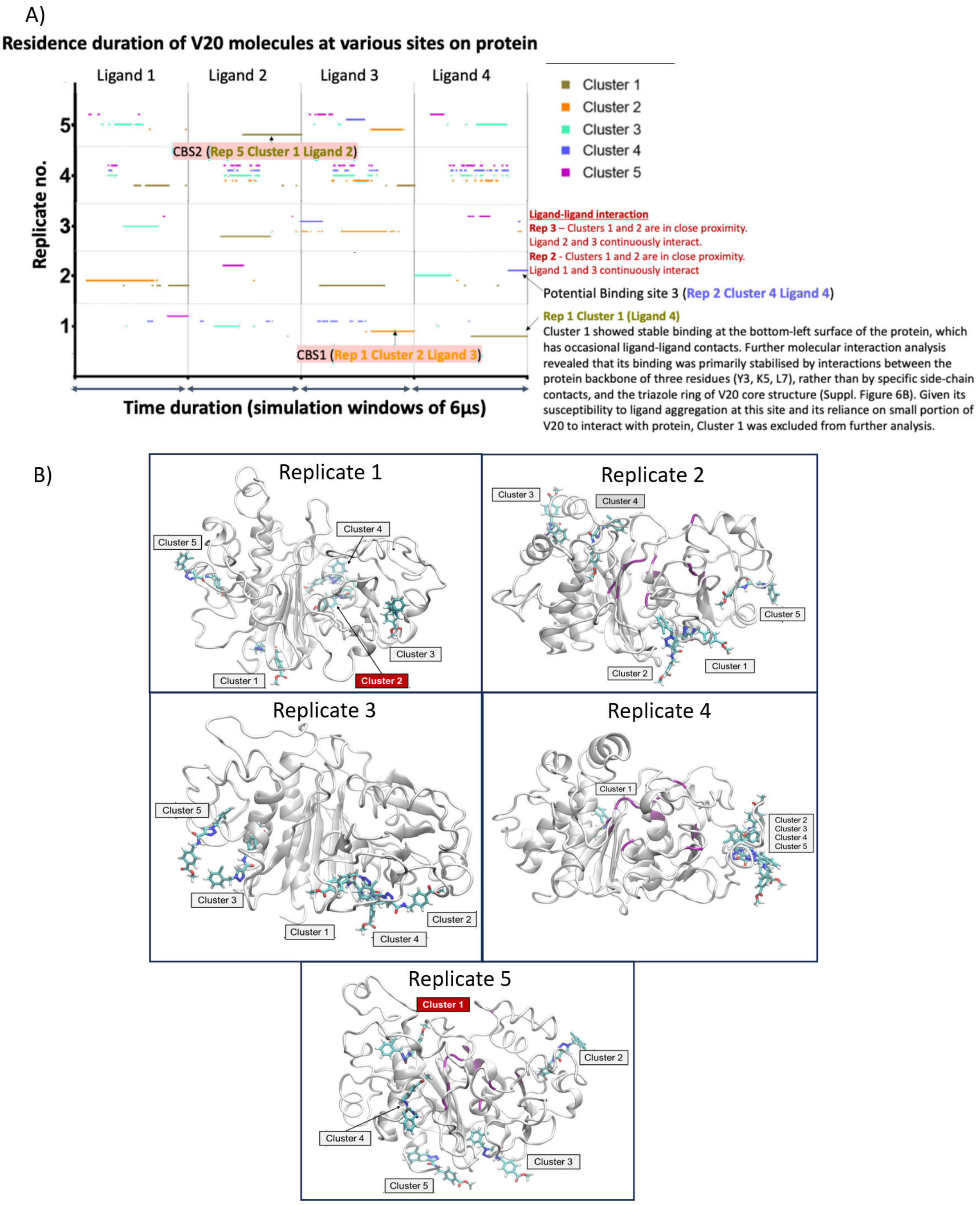


**Fig S6. Molecular dynamics simulations reveal two stable V20 binding clusters on AtSAL1. (A) Cluster analysis of V20 binding across five independent replicates.**
Accelerated molecular dynamics (aMD) simulations with four **V20** inhibitor molecules per replicate, with a total simulation time of approximately 6 µs. The x-axis represents simulation time, divided into four segments corresponding to the four ligands per replicate, while the y-axis separates replicates with black lines. Clusters identified where **V20** maintained a root mean square deviation (RMSD) of ≤5 Å from the cluster centroid. Top five clusters are colour-coded, with Cluster 1 (orange) being the most frequently occupied. Two recurring and stable binding regions, Candidate Binding Site 1 (CBS1) and Candidate Binding Site 2 (CBS2), are annotated. Ligand-ligand interactions are noted in in some replicates.

**(B) Representative views of V20 clusters on AtSAL1 from individual replicates.** Cartoon representations of AtSAL1 protein are shown with bound **V20** molecules from each cluster rendered as licorice sticks and coloured by atom type: nitrogen (blue), oxygen (red), phosphate (khaki), carbon (cyan), chloride (green) and hydrogen (white). Clustered **V20** conformations are representative across the trajectory frames to illustrate binding regions.

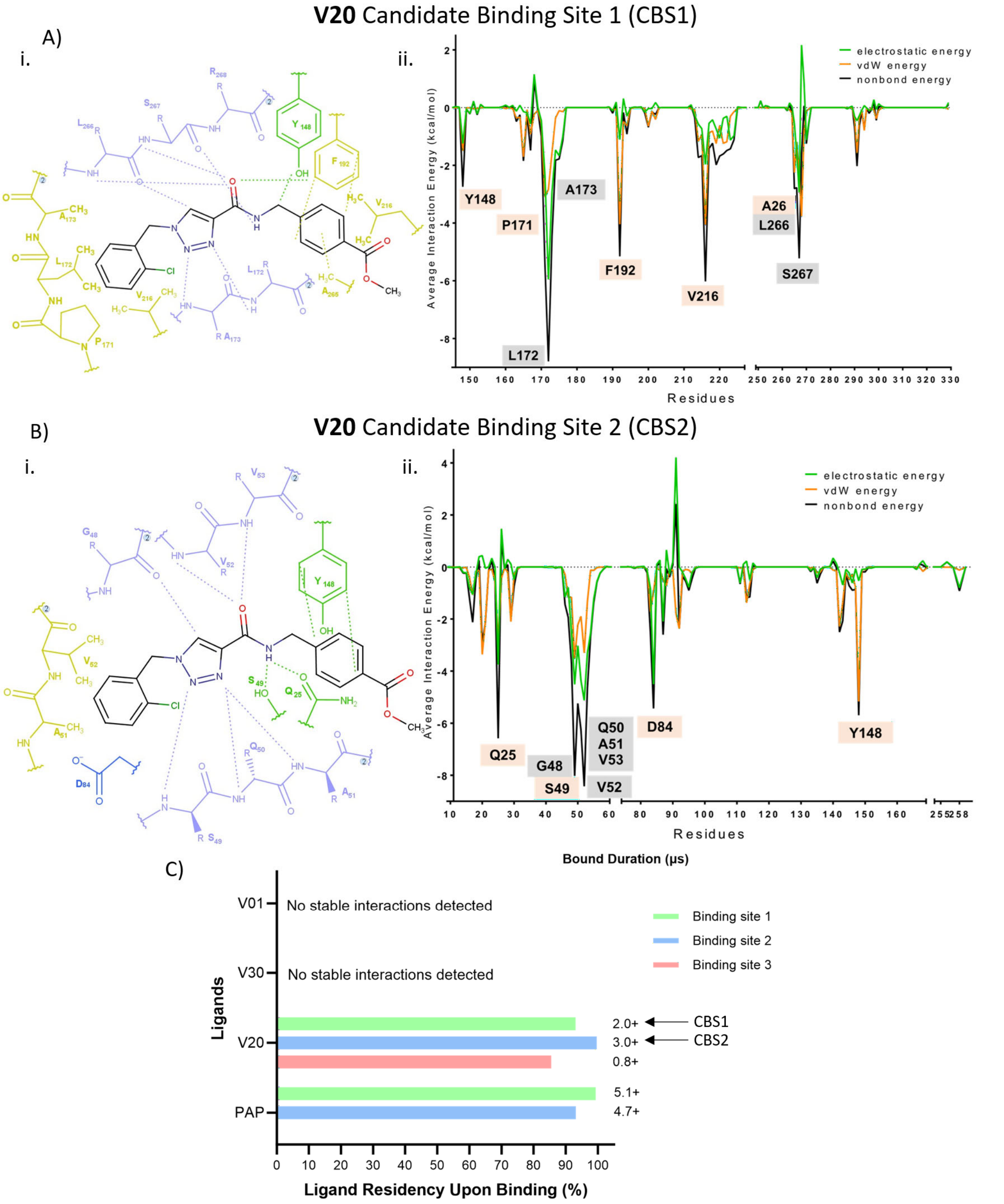


**Fig S7. Interaction energy of AtSAL1 residues with V20 at A) Candidate Binding Site 1 (CBS1) and B) Candidate Binding Site 2 (CBS2).**
**(i) Residue-ligand interaction** diagrams show molecular interactions with residues coloured by coloured by amino acid properties: positively charged (red), negatively charged (blue), polar (green), non-polar (yellow), backbone atoms (purple). Key interaction features between **V20** moieties and surrounding residues are annotated.
**(ii) Interaction energy profiles** at CBS1 (A) and CBS2 (B) show the average nonbonded interaction energies between **V20** and protein residues. Y-axis shows average interaction energy (electrostatic, Van der Waals (vdW) or non-bond energy), with a dotted black line at 0 kcal/mol

distinguishing repulsive (positive) from attractive (negative) interactions. X-axis lists amino acid residues contributing to the interaction. Residues with highest attractive interactions are labelled. **(C) Ligand binding site occupancy and duration.** Bar plot shows of simulation time percentages ligands remained bound at specific sites in aMD simulations. Duration in μs of each interaction is indicated to the right of the bars. A "+" means interaction persisted to simulation end; absence indicates dissociation. Sites where transient ligand-ligand aggregation or non-specific binding occurred were excluded from this analysis.

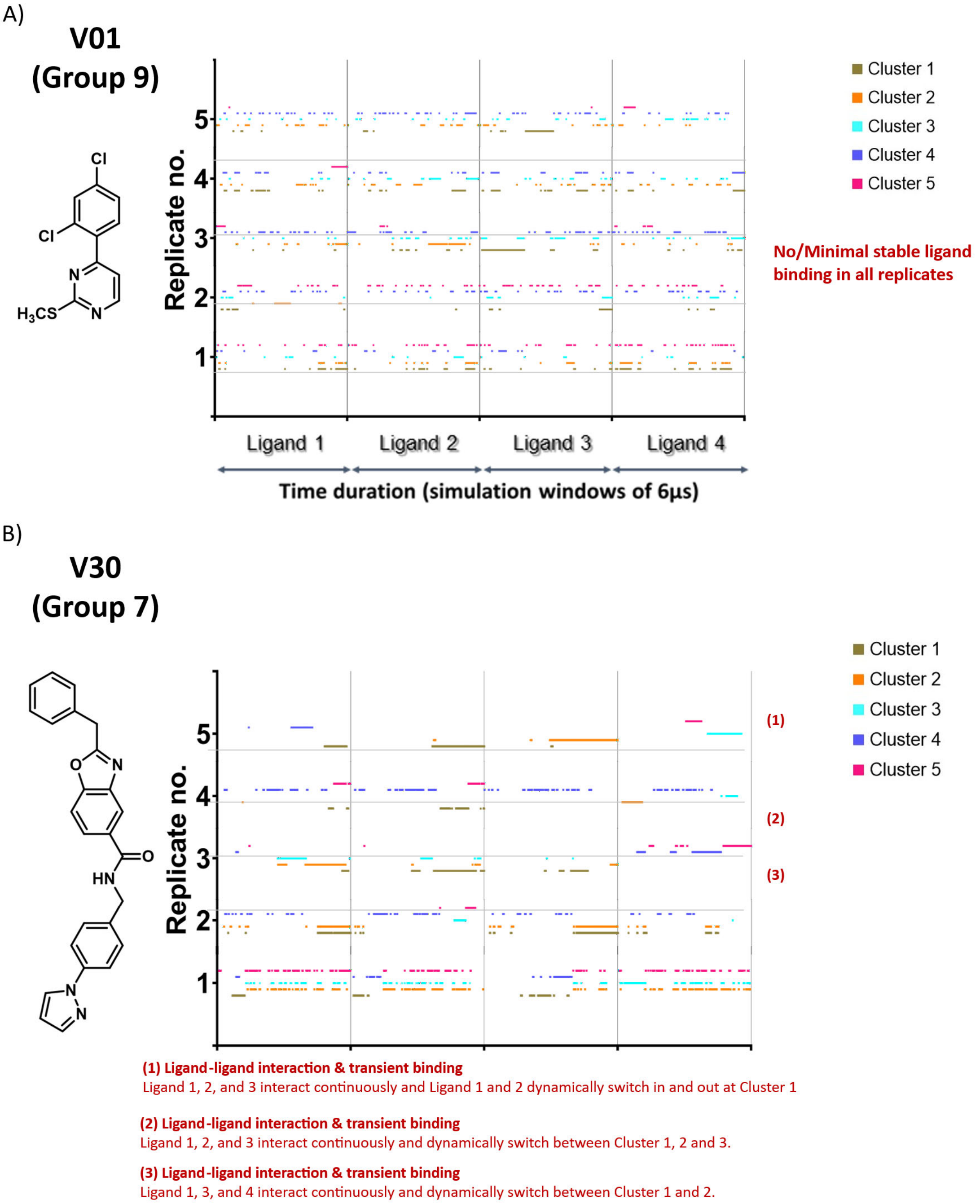


**Fig S8. Cluster analysis of V01 and V30 binding across five independent replicates. (A, B) Binding cluster analysis of inhibitors V01 (A) and V30 (B) on AtSAL1** from five independent replicates, each with four inhibitor molecules (either V01 or V30), with a total simulation time of approximately 6 µs. Trajectory frames were clustered per replicate, and interaction durations were visualised as timelines. Y-axis shows five replicates; x-axis shows simulation time, segmented into four consecutive intervals for four ligands each. Clusters were defined based on trajectory frames in which the ligand maintained a root mean square deviation (RMSD) of ≤5 Å from the centroid structure. Clusters 1 to 5 are colour-coded, with Cluster 1 denoting the most populated binding conformation across simulations.

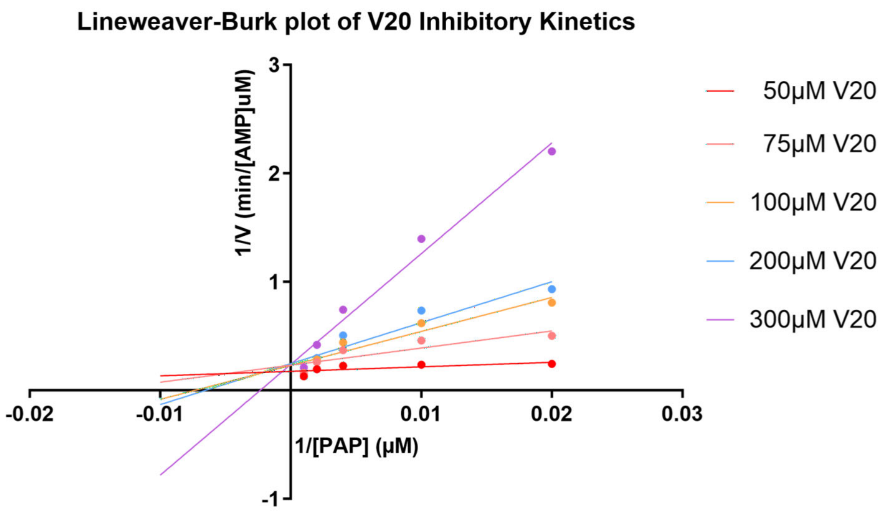


| | 50μM V20 | 75μM V20 | 100μM V20 | 200μM V20 | 300μM V20 |
|---|---|---|---|---|---|
| $K_m$ (μM) | 109.7 ± 54.64<br>a | 448.2 ± 162.1<br>a | 1189.0 ± 394.6<br>a,b | 2729.0 ± 1330.0<br>b | 6606.0 ± 3535.0<br>b |
| $V_{max}$ (μM/min) | 7.4 ± 0.9<br>a | 8.8 ± 1.4<br>a | 13.5 ± 2.8<br>a | 23.5 ± 8.9<br>a | 35.8 ± 17.00<br>a |
| $K_{cat}$ ($min^{-1}$) | 109.9 ± 13.4<br>a | 130.7 ± 20.8<br>a | 200.5 ± 41.6<br>a | 349.0 ± 132.2<br>a | 531.6 ± 252.4<br>a |
| $K_{cat}$/ $K_m$ ($min^{-1}$ $μM^{-1}$) | 1.002 ± 0.514<br>a | 0.292 ± 0.115<br>a | 0.169 ± 0.066<br>a | 0.128 ± 0.079<br>a | 0.0805 ± 0.0576<br>a |

**Fig S9. Determination of AtSAL1 enzyme kinetics in the presence of increasing V20 concentrations.**
Lineweaver-Burk plot shows enzyme kinetics under increasing **V20**. Kinetic parameters ($K_m$, $V_{max}$, $K_{cat}$ and $K_{cat}/K_m$) suggest predominantly competitive inhibition; $K_m$ showing ~60-fold from 50 to 300 μM **V20**; $K_{cat}$ marginally affected (n=3). Two-way ANOVA with Tukey's multiple comparisons test was performed to determine the significance of the kinetic parameters between different **V20** concentrations. a and b ($P < 0.05$) show significant differences between **V20** concentrations.

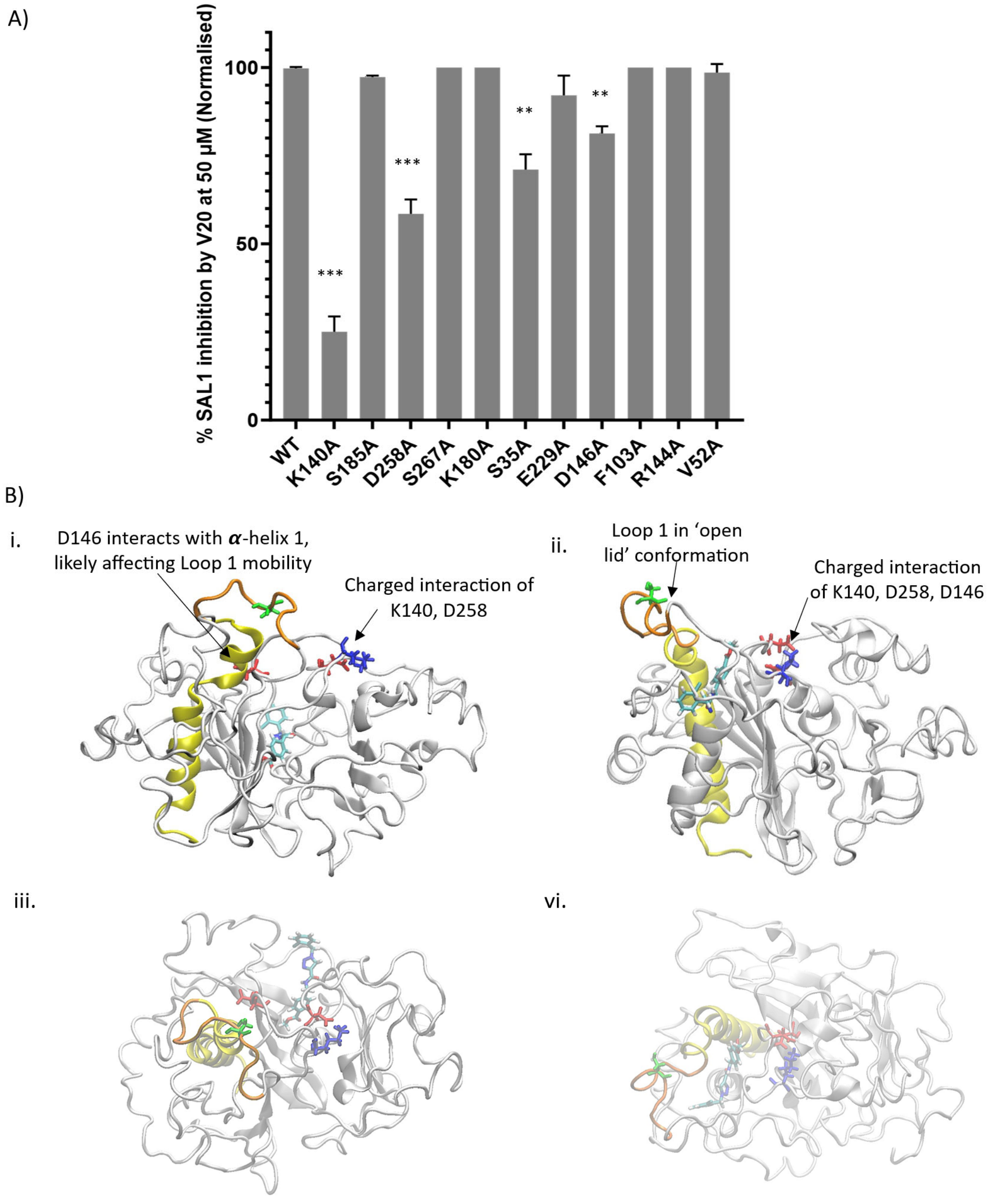


**Fig S10. Inhibitory effect of V20 on AtSAL1 mutants and structural mapping of key residues affecting V20 potency.**

**(A)** Enzymatic activities of wild-type (WT) and selected mutants in 50 µM **V20** and 85 µM PAP substrate (n=3), normalised to each variant's no-inhibitor control. Relative **V20** sensitivity changes shown after baseline subtraction in WT. Asterisks denote statistical significance compared to WT protein inhibition: *P* < 0.05 (*), *P < 0.01 (**), P < 0.001 (***)*.

**(B)** Cartoon representations of AtSAL1 showing residue positions influencing **V20** sensitivity from (i, ii) side-on and (iii, iv) top-down views. Amino acids are colour based on their properties: K140 (blue, positively charged), D258 and D146 (red, negatively charged), and S35 (green, polar). α-helix adjacent to loop 1 is in yellow, and loop 1 in orange. **V20** is in licorice model and coloured by atom type: nitrogen (blue), oxygen (red), phosphate (khaki), carbon (cyan), chloride (green) and hydrogen (white). Arrows indicate charged interactions between residues that may influence local conformations and structural integrity required for catalytic function and inhibitor binding.

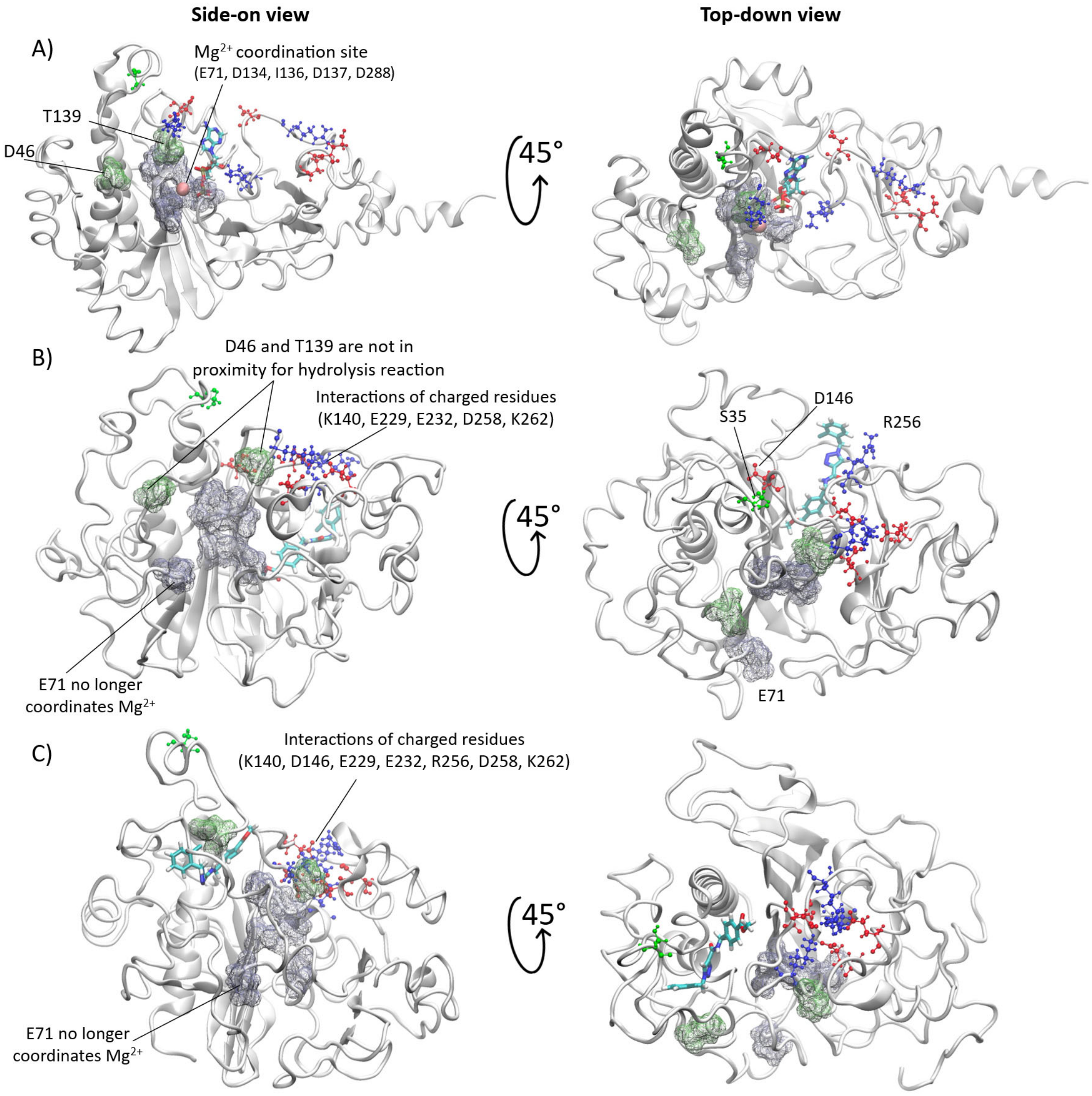


**Fig S11. AtSAL1 catalytic site conformation change upon V20 binding.**
Protein conformational differences in the catalytic site when bound to **(A)** PAP alone, **(B) V20** at CBS1 and **(C)** V20 at CBS2 in side-on (left) and top-down (right) views from accelerated molecular dynamics (aMD) simulations. Key catalytic residues involved in $Mg^{2+}$ and water coordination (E71, D134, I136, D137, D288) are in grey surface representation. Residues directly participating in the hydrolysis reaction (D46, T139) are shown in green surface representation. **V20** is in licorice coloured by atom type: nitrogen (blue), oxygen (red), phosphate (khaki), carbon (cyan), chloride (green) and hydrogen (white). Charged residues engaged in interactions (K140, E229, E232, D258, K262, and D146 and R256 in CBS2) and residues influencing **V20** inhibitory potency (S35, R146, K140, D258) are shown in CPK format, colour-coded by chemical property: positively charged (blue), negatively charged (red), and polar (green). Annotations indicate disrupted coordination or altered residue orientation changes associated with **V20** binding, which may interfere with the catalytic configuration required for PAP hydrolysis.

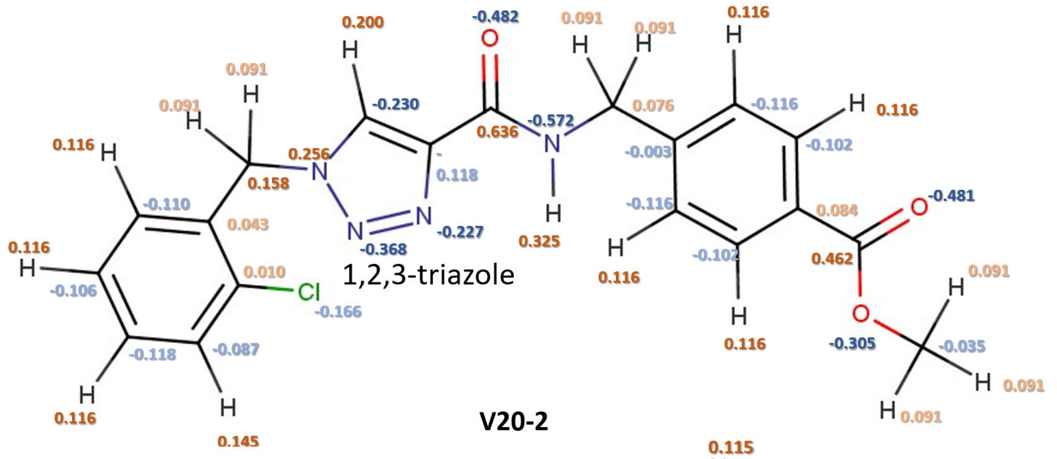


**V20-2**

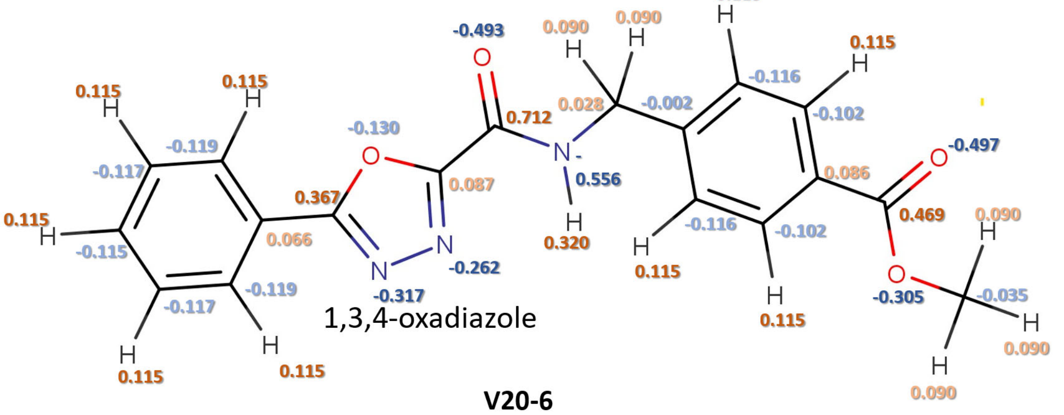


**V20-6**

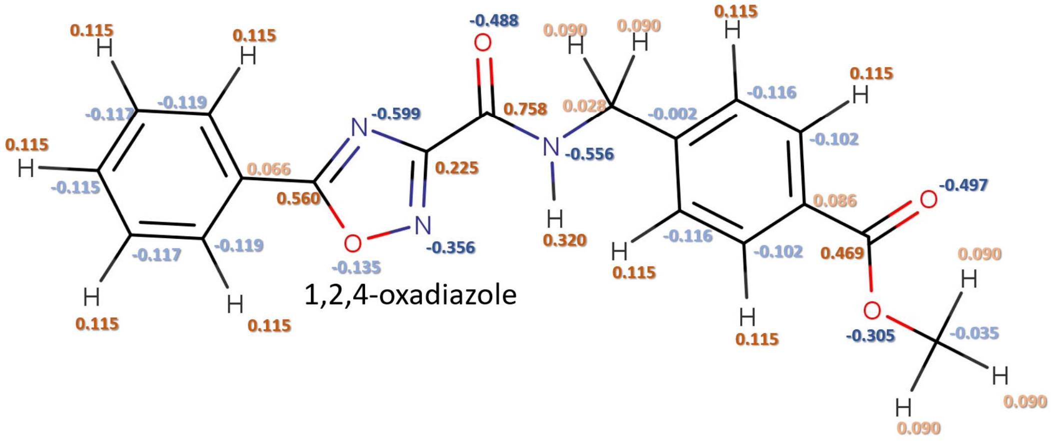


**Fig S12. Chemical structures of V20 and analogues with atomic partial charges.** Structures of the parent compound **V20** and analogues, **V20-2** and **V20-6**, shown with atomic partial charges, computed using CHARMM general force field. Core heterocycles labelled: 1,2,3-triazole (**V20**), 1,3,4-oxadiazole (**V20-2**) and 1,2,4-oxadiazole (**V20-6**). Partial atomic charges are indicated near the atoms: positive charges are shown in shades of orange (light orange for smaller charges, dark orange for larger charges), while negative charges are shown in shades of blue (light blue for smaller charges, dark blue for larger charges). Charge distribution across the triazole/oxadiazole rings, carboxamide, and aromatic groups highlights electronic differences introduced by heterocycle substitution, which may influence hydrogen bonding, ligand binding properties and inhibitory activity.

**Tables**

**AtSAL1 Crystallography Diffraction Data**

| | |
|---|---|
| PDB ID | 8F9Y |
| **Data collection** | |
| Space group | P $6_1$ 2 2 |
| **Cell dimensions** | |
| a, b, c (Å) | 143.0 143.0 75.1 |
| α, β, γ (°) | 90.0 90.0 120.0 |
| Resolution (Å) | 39.72 - 2.60 (2.72 - 2.60) |
| $R_{merge}$ | 0.195 (15.167) |
| $R_{pim}$ | 0.022 (1.682) |
| I/σI | 29.5 (0.8) |
| $CC_{1/2}$ | 1.000 (0.365) |
| Completeness (%) | 100.0 (100.0) |
| Redundancy | 78.2 (81.3) |
| **Refinement** | |
| Resolution (Å) | 39.72 - 2.60 (2.80 - 2.60) |
| No. reflections | 14383 (1401) |
| $R_{work}/R_{free}$ | 0.2254/0.2576 (0.3637/0.3990) |
| No. atoms | 2649 |
| Protein | 2628 |
| Ligand/ion | 11 |
| Water | 10 |
| B-factors (overall) | 128.4 |
| Protein | 128.4 |
| Ligand/ion | 146.7 |
| Water | 118.0 |
| **R.m.s. deviations** | |
| Bond lengths (Å) | 0.003 |
| Bond angles (Å) | 0.56 |

**Table S1. Diffraction data and resolved structure information of AtSAL1 protein.**

Crystallographic data collection, refinement statistics, and geometric parameters for the AtSAL1 protein structure (PDB ID 8F9Y), including unit cell dimensions, resolution range, R-factors, B-factors, and atomic details.

**Primers used in Site-directed Mutagenesis**

| Gene | Primer sequences | |
|---|---|---|
| AtSAL1-D32A | tcaaaaggctttgttgcaatcagctgtgcaatcaaaatctgataaaa | Forward primer |
| | ttttatcagattttgattgcacagctgattgcaacaaagccttttga | Reverse primer |
| AtSAL1-S35A | ctttgttgcaatcagatgtgcaagcaaaatctgataaaagtccagtg | Forward primer |
| | cactggacttttatcagattttgcttgcacatctgattgcaacaaag | Reverse primer |
| AtSAL1-K36A | gttgcaatcagatgtgcaatcagcatctgataaaagtccagtgacc | Forward primer |
| | ggtcactggacttttatcagatgctgattgcacatctgattgcaac | Reverse primer |
| AtSAL1-K140A | tcccctcagaaatcctgcagtgccatcaattggatccaag | Forward primer |
| | cttggatccaattgatggcactgcaggatttctgagggga | Reverse primer |
| AtSAL1-R144A | tggcactaaaggatttctggcgggagatcaatacgcagta | Forward primer |
| | tactgcgtattgatctcccgccagaaatcctttagtgcca | Reverse primer |
| AtSAL1-D146A | aggatttctgaggggagctcaatacgcagtagcac | Forward primer |
| | gtgctactgcgtattgagctcccctcagaaatcct | Reverse primer |
| AtSAL1-E229A | gaggcatcgttcttcgcgtcattcgaaggagct | Forward primer |
| | agctccttcgaatgacgcgaagaacgatgcctc | Reverse primer |
| AtSAL1-D258A | ctccaccagtccgtattgctagccaagcaaagtatg | Forward primer |
| | catactttgcttggctagcaatacggactggtggag | Reverse primer |
| AtSAL1-Y47A | caactgcttgtgaaccagcatcagcaacggtcactggact | Forward primer |
| | agtccagtgaccgttgctgatgctggttcacaagcagttg | Reverse primer |
| AtSAL1-V52A | ctttttctaagactaaactaacagctgcttgtgaaccataatcagca | Forward primer |
| | tgctgattatggttcacaagcagctgttagtttagtcttagaaaaag | Reverse primer |
| AtSAL1-F103A | tagacaaagtagagccattagccgattcctcggtagccaaag | Forward primer |
| | ctttggctaccgaggaatcggctaatggctctactttgtcta | Reverse primer |
| AtSAL1-K180A | cgtctgacgaagatttgttcgcgttgtttcctgctatggatg | Forward primer |
| | catccatagcaggaaacaacgcgaacaaatcttcgtcagacg | Reverse primer |
| AtSAL1-S185A | aggcatccaatttcgtctgccgaagatttgttcttgttg | Forward primer |
| | caacaagaacaaatcttcggcagacgaaattggatgcct | Reverse primer |
| AtSAL1-M202A | tttgaatctaggagctgcgcatatgtccctgaaccaattgtagc | Forward primer |
| | gctacaattggttcagggacatatgcgcagctcctagattcaaa | Reverse primer |
| AtSAL1-S267A | agctccatctcctcttgctaaagctccatactttgc | Forward primer |
| | gcaaagtatggagctttagcaagaggagatggagct | Reverse primer |
| AtSAL1-R268A | agctccatctcctgctgataaagctccatactttgcttgg | Forward primer |
| | ccaagcaaagtatggagctttatcagcaggagatggagct | Reverse primer |

**Table S2. Primer sequences used for site-directed mutagenesis of AtSAL1.**

Primers were designed for PCR-based amplification to introduce single amino acid substitutions in the recombinant AtSAL1 protein. Each primer pair contains the desired nucleotide change for the target residue mutation and was used in a standard site-directed mutagenesis workflow.

**Primers used in RT-qPCR experiments**

| Gene | Primer sequences | Purpose |
|---|---|---|
| AT1G13320 - PROTEIN PHOSPHATASE 2A SUBUNIT A3 (PP2AA3) | F: CATGCAATGGTTACAAGACAAGGTT<br>R: CGAGAAGCGATACTGCACGAA | qPCR internal reference gene |
| Ascorbate Peroxidase 2 (APX2) | F: GCCGTTAGGCTTCTTGACCC<br>R: GGCTCAACTTTGTCCAGTCTACC | PAP-induced stress marker gene |
| Alternative Oxidase 1a (AOX1a) | F: TGGTTGTTCGTGCTGACG<br>R: CACGACCTTGGTAGTGAATATCAG | PAP-induced stress marker gene |
| Calcium Dependent Protein Kinase 32 (CPK32) | F: CTCCCGAGGTGCTAAAACGG<br>R: ACTCCTTGTTCAGTTTCTGCC | PAP-induced stress marker gene |
| Arabidopsis NAC domain containing protein 13 (ANAC013) | F: ACCAGACAGATAAACAATGGATCA<br>R: CAGAAGGAACAGGGTTTAGGAA | PAP-induced stress marker gene |
| Sulfotransferase 12 (SOT12) | F: GGTCACCAATCCACACCTTC<br>R: CGAAATCTGGGGACTCGTAG | PAP-induced stress marker gene |
| Radical Cell Death Induced 1 (RCD1) | F: AGTCAAACCAGGGAGCAAGAGG<br>R: TGGCATCCATGGAGATTTGGG | PAP-induced stress marker gene |

**Table S3. Primers used in RT-qPCR experiments.**

RT-qPCR primer sequences used to quantify expression levels of PAP-responsive genes and a housekeeping gene in Arabidopsis plants treated with **V20** or mock control.
